# Notch dimerization contributes to maintenance of intestinal homeostasis by a mechanism involving HDAC2

**DOI:** 10.1101/2024.04.26.591336

**Authors:** Quanhui Dai, Kristina Preusse, Danni Yu, Rhett A. Kovall, Konrad Thorner, Xinhua Lin, Raphael Kopan

**Affiliations:** State Key Laboratory of Genetic Engineering, School of Life Sciences, Greater Bay Area Institute of Precision Medicine (Guangzhou), Zhongshan Hospital, Fudan University, Shanghai 200438, China; Division of Developmental Biology, Department of Pediatrics, University of Cincinnati College of Medicine and Cincinnati Children’s Hospital Medical Center, Cincinnati, Ohio, USA; Department of Molecular Genetics, Biochemistry and Microbiology, University of Cincinnati College of Medicine, Cincinnati, Ohio, USA

## Abstract

A tri-protein complex containing NICD, RBPj and MAML1 binds DNA as monomer or, cooperatively, as dimers to regulate transcription. Mice expressing Notch <u>d</u>imerization-<u>d</u>eficient (DD) alleles of <u>N</u>otch1 and <u>N</u>otch2 (NDD) are sensitized to environmental insults but otherwise develop and age normally. Transcriptomic analysis of colonic spheroids uncovered no evidence of dimer-dependent target gene miss-regulation, confirmed impaired stem cell maintenance in-vitro, and discovered an elevated signature of epithelial innate immune response to symbionts, the likely underlying cause for heightened sensitivity in NDD mice. TurboID followed by quantitative nano-spray MS/MS mass-spectrometry analyses in a human colon carcinoma cell line expressing either NOTCH2^DD^ or NOTCH2 revealed an unbalanced interactome, with reduced interaction of NOTCH2^DD^ with the transcription machinery but relatively preserved interaction with the HDAC2 interactome suggesting modulation via cooperativity. To ask if HDAC2 activity contributes to Notch loss-of-function phenotypes, we used the HDAC2 inhibitor Valproic acid (VPA) and discovered it could prevent the intestinal consequences of gamma secretase inhibitor (DBZ or DAPT) treatment in mice and spheroids, suggesting synergy between HDAC activity and pro-differentiation program in intestinal stem cells.

## Introduction

The canonical Notch signaling pathway allows adjacent cells to communicate via ligands and receptors embedded in the plasma membrane. Mammals express four Notch receptors: Notch1 (N1), N2, N3 and N4 and 5 ligands: Delta-like1 (Dll1), Dll3, Dll4, Jagged1 (Jag1) and Jag2. Ligand binding to a receptor unfolds a negative regulatory domain to activate a series of consecutive proteolytic cleavages releasing the Notch intracellular domain (NICD). The NICD translocates to the nucleus, where it forms the Notch transcription complex (NTC) together with the co-factors RBPj/CSL (Recombinant binding protein for immunoglobulin J Kappa; CBF1/Suppressor of Hairless/LAG-1) and MAML (Mastermind-like) family members. In the absence of NICD, CSL associates with co-repressors SPEN (*Split Ends* homologue, also called MINT (MSX2 interacting nuclear target) or SHARP (SMRT/HDAC1 associated Repressor Protein)), NCOR1 (nuclear receptor corepressor1), NCOR2 (also called SMART) and others, maintaining transcription in the off-state [1,2]. In the nucleus, the NTCs bind to DNA to regulate target gene expression [3–7]. Binding to the DNA occurs on conserved CSL sites. NICDs can form NTC homo- or hetero-dimers on sequence-paired-sites (SPS) which consists of two CSL sites organized in head-to-head orientation 15-18 nucleotides apart [8–13]. To investigate the biological function of NTC dimers we generated mice with ARG to ALA substitutions at the dimer interface in both Notch1 and Notch2 (henceforth NotchRA or NDD) to block dimerization (N1^R1974A^ or N1^RA^ and N2^R1934A^ or N2^RA^). These mice showed defects in multiple organs, including spleen, heart and the intestine [14]. They proved to be hypersensitive to environmental conditions, with hypo-allergenic environment ameliorating many of the more severe phenotypes (further analyzed in accompanied study - Preusse et al.).

The mammalian intestine, divided into the small intestine and the large intestine or colon, consists of the inner layer of consistently renewing epithelial cells, or intestinal mucosa, surrounded by an outer, innervated smooth muscle layer responsible for peristaltic movement. The epithelium contains adsorptive cells (enterocytes) as well as sensory (taft) and secretory cells (mucus-secreting goblet cells, Paneth cells and enteroendocrine cells). Intestinal stem cells (ISC) reside at the bottom of the crypts [15,16], marked by the expression of leucine rich repeat containing G protein-coupled receptor 5 (Lgr5), achaete-scute family bHLH transcription factor 2 (Ascl2) and olfactomedin 4 (Olfm4) [17–20]. Notch signaling maintains the stem cell niche and regulates differentiation of intestinal stem cells [21–23]. Blocking Notch signaling leads to loss of the stem cell niche and to goblet cell metaplasia [22,24–26]. Consistently, Notch activation expends the stem cell niche at the expense of goblet and enteroendocrine cells [23]. <u>N</u>otch <u>d</u>imerization-<u>d</u>eficient (NDD) mice show an environmentally modulated susceptibility to DSS and a decrease in colonic stem cells [14], but the mechanism linking Notch dimerization with the stem cell niche remains unknown.

In this study we used our NDD mouse model to further investigate the role of NTC dimerization in intestinal and colonic stem cells. To that end we examined the small intestine *in vivo*, as well as colonic and small intestinal spheroids *ex vivo*, to analyze maintenance, differentiation, and proliferation with or without input from the immune system. RNA sequencing of colonic spheroids confirmed a decrease in undifferentiated stem cells and enhanced differentiation accompanied by elevated innate immune response to symbionts but did not show any miss-regulation of dimer-dependent target genes. To identify changes in dimer vs monomer Notch interactome we performed TurboID and quantitative nano-spray MS/MS mass-spectrometry analyses in a human colon carcinoma cell line, HT-29. The constitutively active TurboID-fused Notch2 and its Notch2RA counterpart constructs expressed at similar levels and biotinylated NTC and other known partners but notably, Notch2RA labeling of endogenous human NOTCH2 was significantly reduced relative to control. Qualitatively, the interactome was unchanged; on average, however, quantitative analysis revealed a 33%-39% labeling reduction across all interacting partners, consistent with weak (hypomorphic) activity of NotchRA. Interestingly, the HDAC2 interactome was significantly less affected by loss of dimerization. We hypothesize that dimer-deficiency and loss of cooperativity altered the balance between transcription machinery interactions vs. the dimer independent negative regulatory/deacetylase complex, somehow sensitizing the animal to environment-mediated insults, and perhaps regulating the level of protein acetylation among interacting partners in a dimer independent fashion which might explain the relatively mild phenotype in the absence of microbiome [14]. To test the assumption that HDAC2 activity contributes to Notch loss of function phenotypes we used the HDAC2 inhibitor Valproic acid (VPA) and discovered it could block the intestinal consequences of gamma secretase inhibitor treatment (DBZ or DAPT) in mice and spheroids, respectively, suggesting synergy between HDAC activity and pro-differentiation program in intestinal stem cells.

## Results

### Notch dimerization-deficiency compromises colonic and intestinal spheroids formation

In a previous study we reported that *N1^RA/RA^; N2^RA/RA^* mice loose colonic stem cells when challenged with 1% DSS, a dose that is well tolerated by wild type controls [14]. To test if loss of Notch dimerization affects colonic stem cells via a cell-autonomous mechanism, we established colon spheroid cultures devoid of immune system components and passaged them *in vitro.* We observed that *N1^RA/RA^; N2^RA/RA^* cells generated the fewest number of spheroids (average of 2-3 per field), *N1^+/+^; N2^RA/RA^* cells generated an intermediate number (∼6/field) and *N1^+/+^; N2^+/+^* cells generated the most (∼15 per field; Fig 1A and 1B). To assess the fraction of stem cells in colon spheroids, we performed qPCR analyses of *Lgr5* levels on mRNA collected from three individual experiments of passage 1 spheroids. This assay confirmed that *N1^RA/RA^; N2^RA/RA^* spheroids had fewer *Lgr5*+ cells compared to wild type controls (Fig 1C). Accordingly, dissociating the spheroids into cell suspension and embedding the same number of cells in Matrigel (passage 2) again showed fewer spheroids derived from *N1^RA/RA^; N2^RA/RA^,* more from *N1^+/+^; N2^RA/RA^* cells but still fewer than *N1^+/+^; N2^+/+^* control (Fig 1D). We quantified cell numbers instead of spheroid count in passage 1 and 2 to illustrate the deficiency of self-renewal in colonic stem cells lacking a full complement of dimerization-competent Notch receptors (Fig 1E). Consistently, baseline *in vivo Lgr5* expression is reduced in the distal colon of *N1^RA/RA^; N2^RA/RA^* mice compared to wild type controls and was further reduced after treatment with 1% DSS (S1 Fig A). Orthogonal support for stem cell stress was provided by analyzing *N1^RA/RA^; N2^RA/RA^; Lgr5-EGFP-IRES-creERT2* mice, which lost weight rapidly from day 4 relative to *N1^+/+^; N2^+/+^; Lgr5-EGFP-IRES-creERT2* when exposed to 2% DSS to induce colitis (S1 Fig B). GFP staining confirmed a reduction in stem cell numbers in the colons of NDD mice after DSS treatment (S1 Fig C and D).

**Fig 1.**
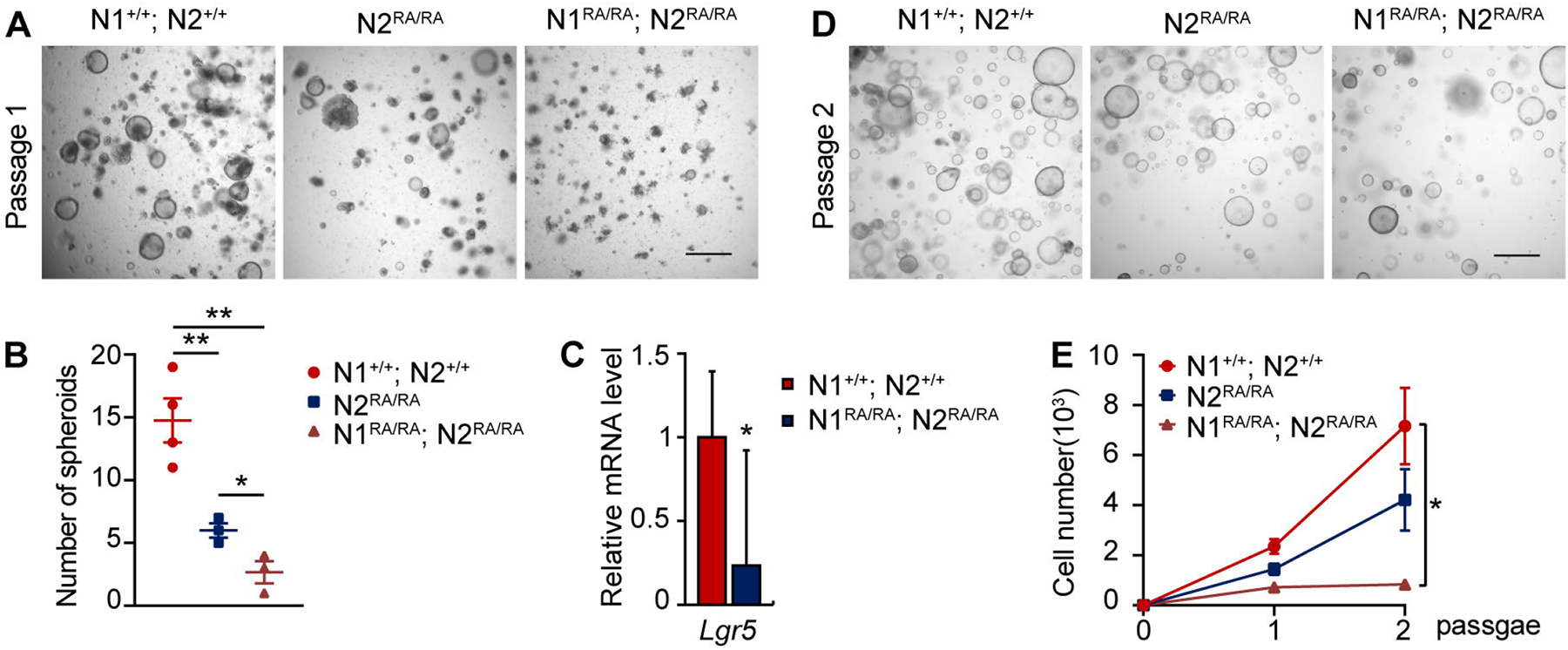
Notch dimer-deficiency compromises colon spheroid formation. **A**. Representative bright-field images of passage 1 colon spheroids established from isolated adult mice colon crypts. Scale bars=500 μm. Dark, opaque clusters are cellular debris. **B**. The number of spheroids present at passage 1, average of 3 fields in at least 3 independent experiments. **C**. qPCR analysis of gene expression of *Lgr5* on RNA extracted from passage 1 colon spheroids. n=3 per group. **D**. Colon spheroids 4 days after passage, representative bright-field images. **E**. Quantification of cell numbers (single cell and clumps) at indicated passage. Quantitative data are presented as mean ± SEM. \**p* <0.05, \*\**p* <0.01, \*\*\**p* <0.001.

NICD accumulation is graded along the A/P axis of the intestine, with the lowest amount detected in the colon (Supplemental Figure S3 in [25]). To investigate if this gradient was reflected in greater sensitivity to Notch dimerization in the colon, we asked if anterior small intestine spheroids will be affected to the same degree as the posterior ones by loss of dimerization. As we observed in the colon, smaller spheroids with fewer “crypts” were derived from *N1^RA/RA^; N2^RA/RA^* mice relative to those derived from the wild type controls (Fig 2A, asterisk). In addition, fewer Olfm4 positive stem cells and lower levels of *Olfm4* mRNA were detected in *N1^RA/RA^; N2^RA/RA^* spheroids, which also have fewer Ki67 positive cells indicating decreased proliferation potential and reduced stem cell numbers (Fig 2B, 2C). Reduction in stem cells in *N1^RA/RA^; N2^RA/RA^* spheroids was accompanied by accumulation of lysozyme-positive Paneth cells, suggesting a defect in self renewal leading to enhanced differentiation (Fig 2D).

**Fig 2.**
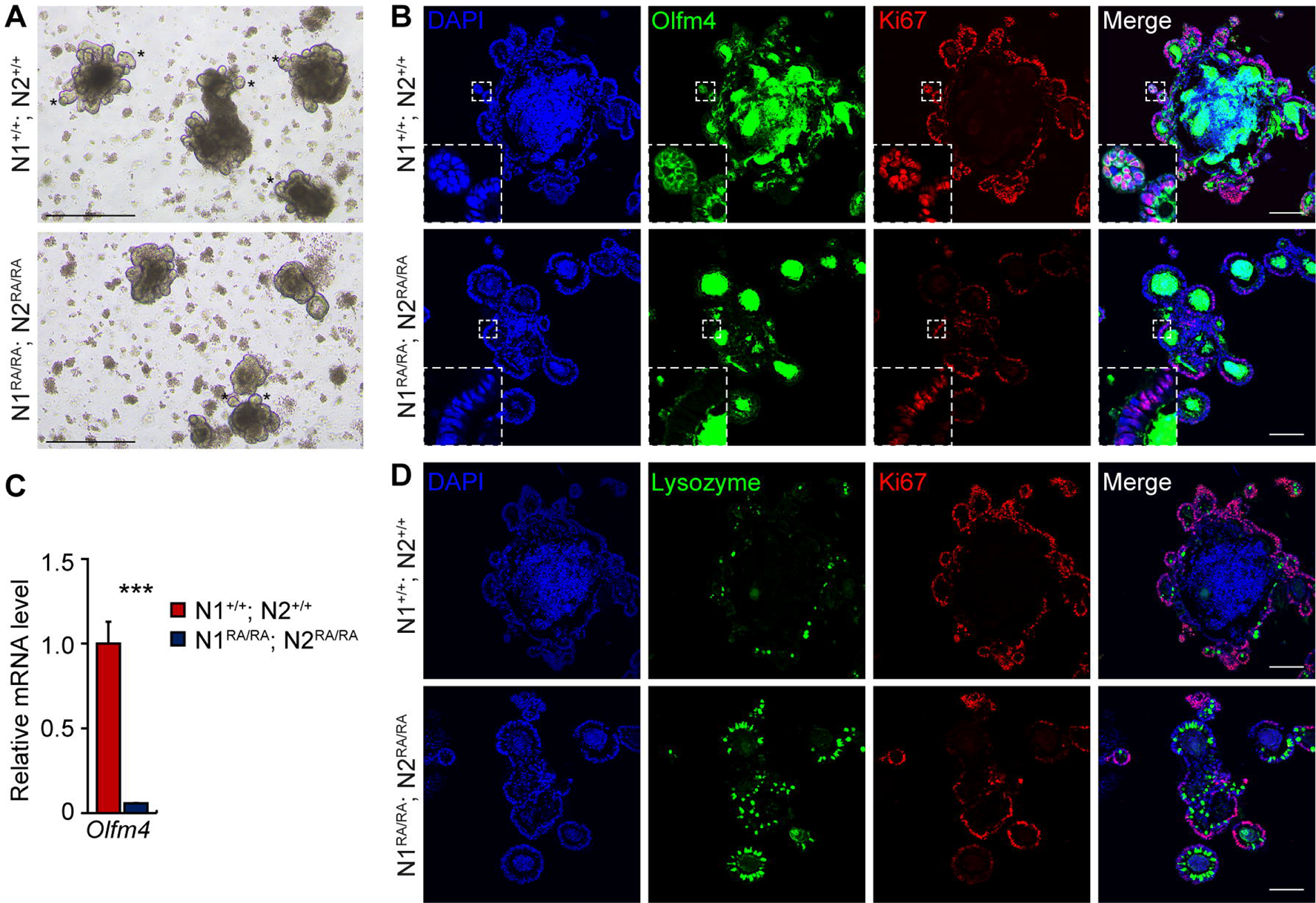
Notch dimer-deficiency compromises small intestinal spheroid formation. **A**. Representative bright-field images of small intestine spheroids at day 6 of passage 1. Asterisk marks examples of crypt. Scale bars=500 μm. **B**. Immunofluorescence staining of Olfm4 (green) and Ki67 (red) in small intestinal spheroids, nuclei staining with DAPI. White dashed box indicates regions shown in higher magnifications. Scale bars=100 μm. **C**. qRT-PCR analysis of gene expression of *Olfm4* in spheroids. Quantitative data is presented as mean ± SEM. \**p* <0.05, \*\**p* <0.01, \*\*\**p* <0.001. **D**. Immunofluorescence staining of Lysozyme (Paneth cells, green) and Ki67 (red) in spheroids, nuclei staining with DAPI. Scale bars=100 μm. n=3 per group.

### Loss of Olfm4^+^ stem cells and increase of Paneth cells in small intestine of Notch dimer-deficient mice

To test the hypothesis that the RA mutants are a mild loss of function allele, we compared the small intestine in *N1^RA/RA^; N2^RA/RA^* and wild type mice in which Notch signaling was removed or inhibited. Loss of Notch signaling by inhibition (with DBZ) or with N1 and N2 neutralizing antibodies leads to loss of stem cells and enhanced differentiation towards secretory cell lineages [20]. We analyzed intestinal tissue collected from adult *N1^RA/RA^; N2^RA/RA^* and control mice for expression of the stem cell markers *Lgr5*, *Ascl2* and *Olfm4* using qRT-PCR transcript analysis and immunofluorescence for the protein products. The expression level of *Lgr5* and *Ascl2* in the small intestine was unaffected in *N1^RA/RA^; N2^RA/RA^* (S3 Fig A). By contrast, *N1^RA/RA^; N2^RA/RA^* mice showed a marked decrease in numbers of cells positive for Olfm4 (Fig 3A), consistent with a significant decrease of *Olfm4* mRNA abundance (Fig 5B). The expression of another progenitor cell marker, *Prom1,* in *N1^+/+^; N2^+/+^* (WT) and *N1^RA/RA^; N2^RA/RA^* mice was indistinguishable (S3 Fig A), as was Sox9 staining in jejunum and colon (S3 Fig B and C). Likewise, whereas inhibition of Notch signaling leads to elevated Atoh1 expression, *Atoh1* levels in the *N1^RA/RA^; N2^RA/RA^* jejunum and ileum remained indistinguishable from wild type controls (Fig 3C), indicating sufficient levels of Notch activity. To determine the functional significance of reduced Olfm4^+^ stem cell numbers in *N1^RA/RA^; N2^RA/RA^* mice, we tested for cellular proliferation and apoptotic cell death by Ki67, PCNA and cleaved Caspase-3 staining. *N1^RA/RA^; N2^RA/RA^* small intestine had a reduced *Ki67* mRNA abundance and PCNA staining (Fig 3A and 3B), but cleaved caspase-3 was not elevated (S3 Fig D). Loss of Notch dimerization displayed one consequence of Notch signaling inhibition, namely, a robust increase in lysozyme-positive and MMP7-positive Paneth cells (Fig 3D and S3 Fig E). However, Enteroendocrine (CgA), Tuft (Dcamkl1), and Goblet cell (Muc2) numbers were not affected by loss of Notch dimerization (S3 Fig F-H).

**Fig 3.**
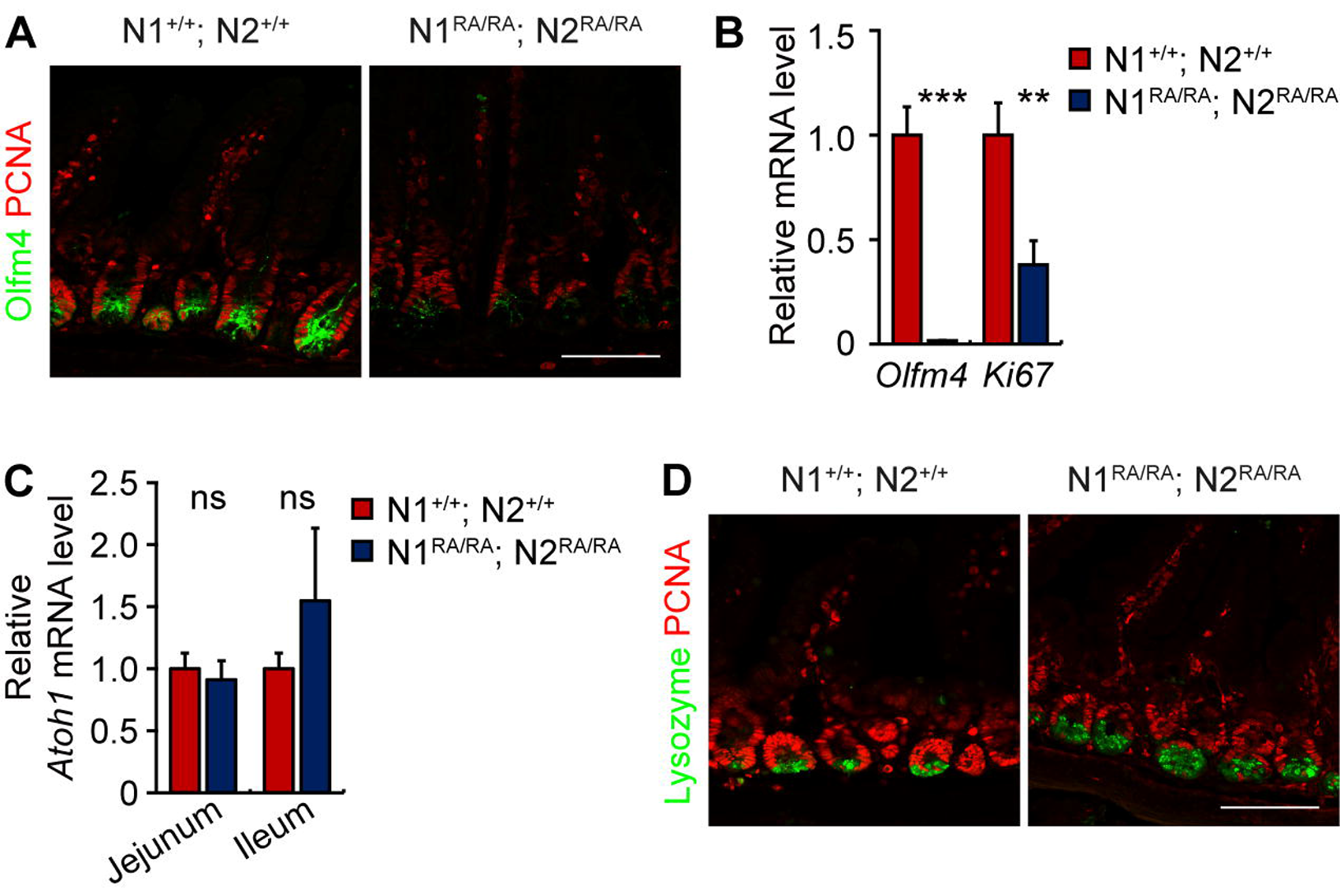
Loss of Olfm4^+^ stem cells and increased Paneth cells in small intestine of NDD mice. **A**. Representative images of Olfm4 and PCNA immunofluorescence staining of jejunum from *N1^+/+^; N2^+/+^* and *N1^RA/RA^; N2^RA/RA^* mice. n=3 mice per group. Scale bars=100 μm. **B**. qRT-PCR analysis of gene expression of stem cell marker *Olfm4* and *Ki67*. Quantitative data are presented as mean ± SEM. \**p* <0.05, \*\**p* <0.01, \*\*\**p* <0.001. **C**. qRT-PCR analysis of *Atoh1* gene expression in jejunum and ileum. n=3 mice per group. Scale bars=100 μm. Quantitative data are presented as mean ± SEM. ns-Not significant. **D**. Lysozyme and PCNA immunofluorescence staining of jejunum from *N1^+/+^; N2^+/+^* and *N1^RA/RA^; N2^RA/RA^* mice. n=3 mice per group. Scale bars=100 μm.

Collectively, these data suggest that *Olfm4* gene expression was sensitive to Notch dimerization or that it required the full strength of Notch signal. Accelerated Paneth cell differentiation without Atoh1 upregulation is another hallmark of a weak loss of function Notch allele, perhaps acting as sentinels for reduced Notch activity in stem cells.

### N2 plays a greater role than N1 in protecting the colon from DSS

To assess the susceptibility of mice with single dimerization deficient Notch to DSS-induced colitis, we subjected *N1^RA/RA^*; *N2^+/+^, N1^+/+^; N2^RA/RA^*, and *N1^RA/RA^; N2^RA/RA^* mice as well as wild type controls to two 10-day cycles of 1% DSS treatment, with a 14-day normal drinking water recovery phase. During the initial DSS cycle, all NDD mice lost weight, which was recovered after removal of DSS from drinking water. During the second DSS cycle, however, WT and *N1^RA/RA^*; *N2^+/+^* mice showed only minor weight reductions, whereas *N1^+/+^; N2^RA/RA^* and *N1^RA/RA^; N2^RA/RA^* mice suffered a more pronounced and rapid decline in body weight (Fig 4A). Macroscopic and histological analyses detected no significant disparities in colon length among the experimental groups at baseline, and the colon appeared normal in all mice (Fig 4B and 4C). However, after exposure to 1% DSS, the colon of *N1^RA/RA^*; *N2^+/+^, N1^+/+^; N2^RA/RA^*, and *N1^RA/RA^; N2^RA/RA^*was notably shorter than that of controls – consistent with enhanced inflammation (Fig 4B). Further, *N1^+/+^; N2^RA/RA^* and *N1^RA/RA^; N2^RA/RA^* mice exhibit more pronounced histological injury compared to both WT and *N1^RA/RA^*; *N2^+/+^* mice (Fig 4C). Immunofluorescence analysis using Ki67 staining demonstrated decreased cellular proliferation in *N1^+/+^; N2^RA/RA^* and *N1^RA/RA^; N2^RA/RA^* mice subjected to 1% DSS treatment, but no significant differences under baseline conditions (Fig 4D).

**Fig 4.**
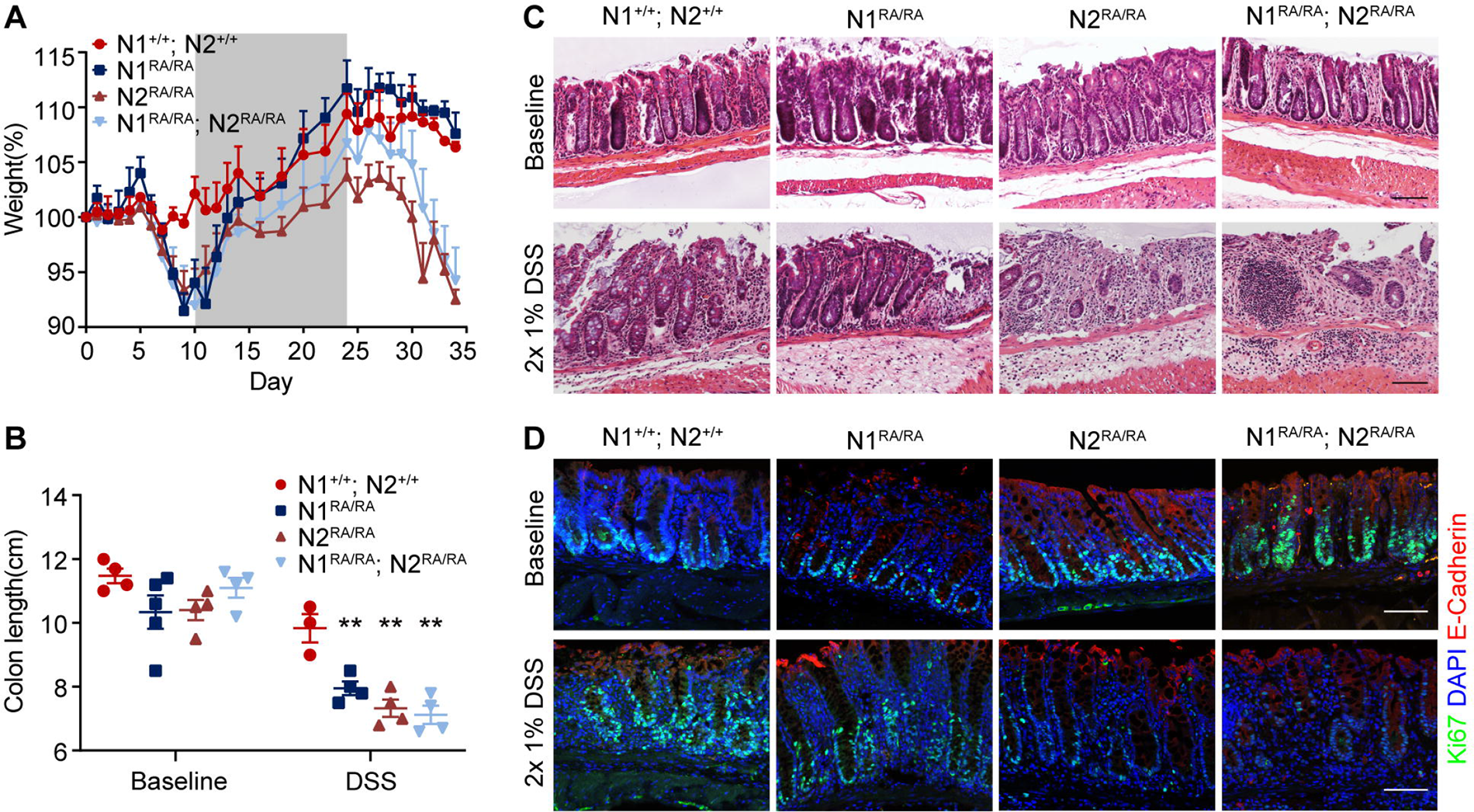
N2 plays a greater role than N1 in protecting the colon from DSS. **A**. Daily weight measurements of *N1^+/+^; N2^+/+^*, *N1^RA/RA^*, *N2 ^RA/RA^*or *N1 ^RA/RA^; N2^RA/RA^*mice treated with alternate 10-day cycles of 1% DSS (white sections) and 14-days without DSS (gray sections). **B**. Colon length measurements of *N1^+/+^; N2^+/+^*, *N1^RA/RA^*, *N2^RA/RA^* or *N1^RA/RA^; N2^RA/RA^* mice treated with 1% DSS or no DSS. Quantitative data are presented as mean ± SEM. \**p* <0.05, \*\**p* <0.01, \*\*\**p* <0.001. **C**. H&E staining of distal colon tissue from DSS-treated or no DSS mice with the indicated genotypes. Scale bars=100 μm. n=3 mice per group. . . Immunofluorescence staining of Ki67 (green), E-cadherin (red) and DAPI (blue) in distal colon tissue from DSS-treated and untreated mice with the indicated genotypes. Scale bars=100 μm. n=3 mice per group.

Despite the reduction in stem cells during DSS exposure, NDD mice recover from the treatment [27]. To test if stem cell function diminished with age, we collected tissue from mice of 300 days of age or older. H&E staining and immunofluorescence imaging of colon sections showed normal histology and numbers of proliferating Ki67^+^ cells in *N1^RA/RA^, N2^RA/RA^* and *N1^+/–^; N2^+/–^* mice (S4 Fig A-C and S4 Fig A’-C’). Together, these results demonstrate that Notch dimerization-deficient intestinal stem cells can maintain homeostasis when unchallenged but are sensitized to exogenous challenges.

### Transcriptomic analysis identifies elevated innate immune signature but no bias against dimer-dependent gene expression

To identify the transcriptomic consequences of losing cooperativity [28], differential gene expression and pathway enrichment analyses were performed on RNA isolated from colonic spheroids (Fig 5, <u>GSE262485</u>) which revealed several interesting consequences of losing dimerization. Unsupervised clustering of transcripts largely divided the 507 differentially expressed genes (DEGs) into up and down regulated genes. Upregulated DEGs show increase in secretory differentiation genes, defense response to symbionts (in a sterile environment devoid of any), and increased epithelialization. Down regulated DEGs confirmed a downregulation of *Lgr5* in NDD mice and show reduced proliferation and impaired adhesion/barrier (S1 Table - Tabs 1-3). Importantly, the DEG analysis was not consistent with the hypothesis that dimerization was only required for targets regulated by SPS sites [29,30], present in over 20% of enhancers based on studies in a T-ALL cell line [31]: expression of dimer-dependent targets such as *NRARP*, *Hes5*, or *Hes1* [29] was unchanged, suggesting that differentially expressed genes reflected an overall reduction in Notch transcriptional activity rather than selective reduction of dimer-dependent gene expression (Fig 5, S1 Table - Tab1). Clustering of putative gene products in String (https://string-db.org) and performing pathway analysis in ToppFun (https://toppgene.cchmc.org/enrichment.jsp) with the FDR cutoff of 0.05 showed that spheroids activated a network of innate immunity defense against symbionts that includes interferon β-dependent antiviral genes (Fig 5 - clusters 3 and 4, S1 Table - tab1 and tab4). A small TEAD4-linked cluster involved in steroid hormone metabolism was present in cluster 2 but did not pass the ToppFun FDR cutoff. Together, these results demonstrate that Notch dimerization is required throughout the A/P axis of the intestine, and that reduced proliferation and enhanced differentiation was coupled with an abnormally elevated state of innate immunity genes explaining in part the adverse response to stripping the mucosa with 1% DSS. It also explains in part the normal in-utero intestinal development is followed by rapid postnatal demise and GI disintegration after colonization by symbionts in *N1^RA/–^; N2^RA/–^* (see [14] and accompanying paper – Preusse et al.).

**Fig 5.**
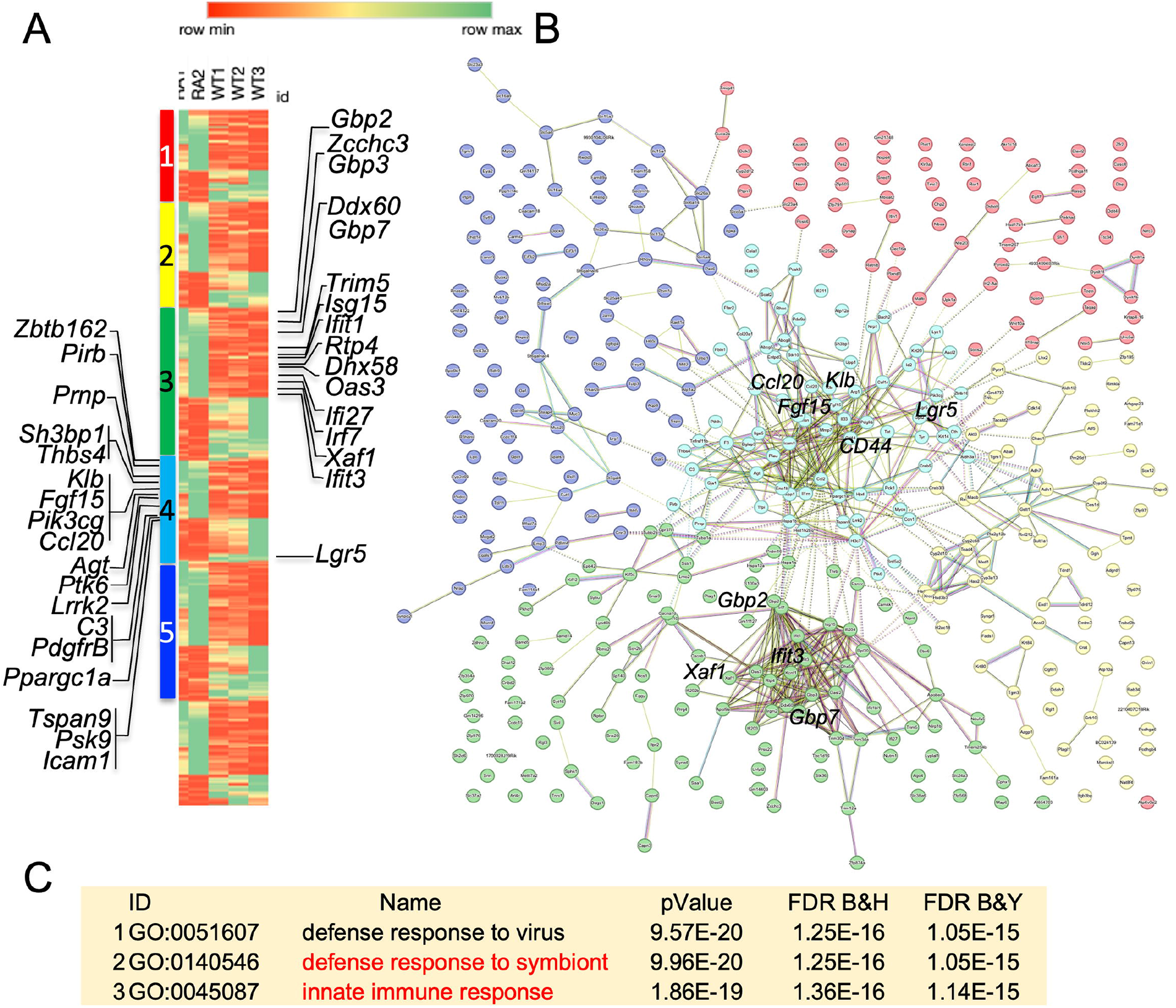
Colon spheroids RNA-seq and cluster analysis. A. mRNA differential expression heatmap sorted by K-means cluster analysis of protein products. WT represents independent repeats from *N1^+/+^; N2^+/+^* controls, RA represents repeats from *N1^RA/RA^; N2^RA/RA^* colon spheroids. Cluster 3 and 4 contain innate immune network proteins, shown with relative position in the heatmap. B. String analysis of protein products. Only cluster 3,4,5 have met the 0.05 FDR cutoff. See Supplemental Table S1, Tab 4 for more information. This cluster map was added to supplemental data as an SVG file to facilitate browsing. C. Pathway analysis of the 33 proteins listed here and 15 listed in Supplemental Table S1, Tab4.

### Loss of dimerization dampens protein-protein interactions of NOTCH

The transcriptome analyses suggested a reduction in adhesion, increased spheroid differentiation, and elevated innate immune apparatus in spheroids, but failed to provide mechanistic insight linking Notch dimerization to these responses. To investigate the hypothesis that dimerization is involved in modulating Notch dimer-dependent protein-protein interactions, we performed TurboID analysis [32,33]. We first established stable HT-29 human colorectal carcinoma cell line overexpressing truncated human NOTCH2^ΔE^-TurboID, NOTCH2^ΔE^RA-TurboID or TurboID as control (henceforth, NICD2-TurboID or NICD2RA-TurboID, selected based on the experiments described in Fig 5). After supplementing the biotin-low DMEM culture medium with 500 μM biotin for one hour, protein samples were harvested from the control, NICD2-TurboID and NICD2RA-TurboID stable polyclonal lines. Biotinylated proteins were captured on streptavidin beads and washed. Western blot analysis confirmed that biotinylated proteins from control, NICD2-TurboID and NICD2RA-TurboID were captured (Fig 6A). The western blot demonstrated that NICD2-TurboID and NICD2RA-TurboID were present in equivalent amounts (Fig 6A and 6B, numbers normalized to endogenous Notch2 from TurboID transfected cells). Protein biotinylation not present in the parental cells depends on self-labeling or on interaction between NICD2RA-TurboID molecules. As expected, streptavidin recovered NICD2-TurboID and endogenous NICD2 proteins far more efficiently from NICD2-TurboID cells than from NICD2RA-TurboID expressing cells, indicating as expected that dimerization increased the efficiency of labeling of the endogenous partner NICD2 (Fig 6A and 6B). Purified biotinylated protein samples were submitted to Beijing Genomics Institute (BGI) for next generation label-free quantitative proteomic analyses on a Q-Exactive HF X (Thermo Fisher Scientific, San Jose, CA) in data independent acquisition (DIA) mode. The ionized peptide masses were aligned against the human proteome. Initial assessments of data quality, including unique peptide number, mass distribution, and protein coverage indicated the labeling met or exceeded accepted thresholds (Fig 6C). Overall, the vast majority of biotinylated proteins were nuclear and linked to transcription. >98% were recovered less efficiently by NICD2RA-TurboID compared to NICD2-TurboID. The full data set is presented in S2 Table.

**Fig 6.**
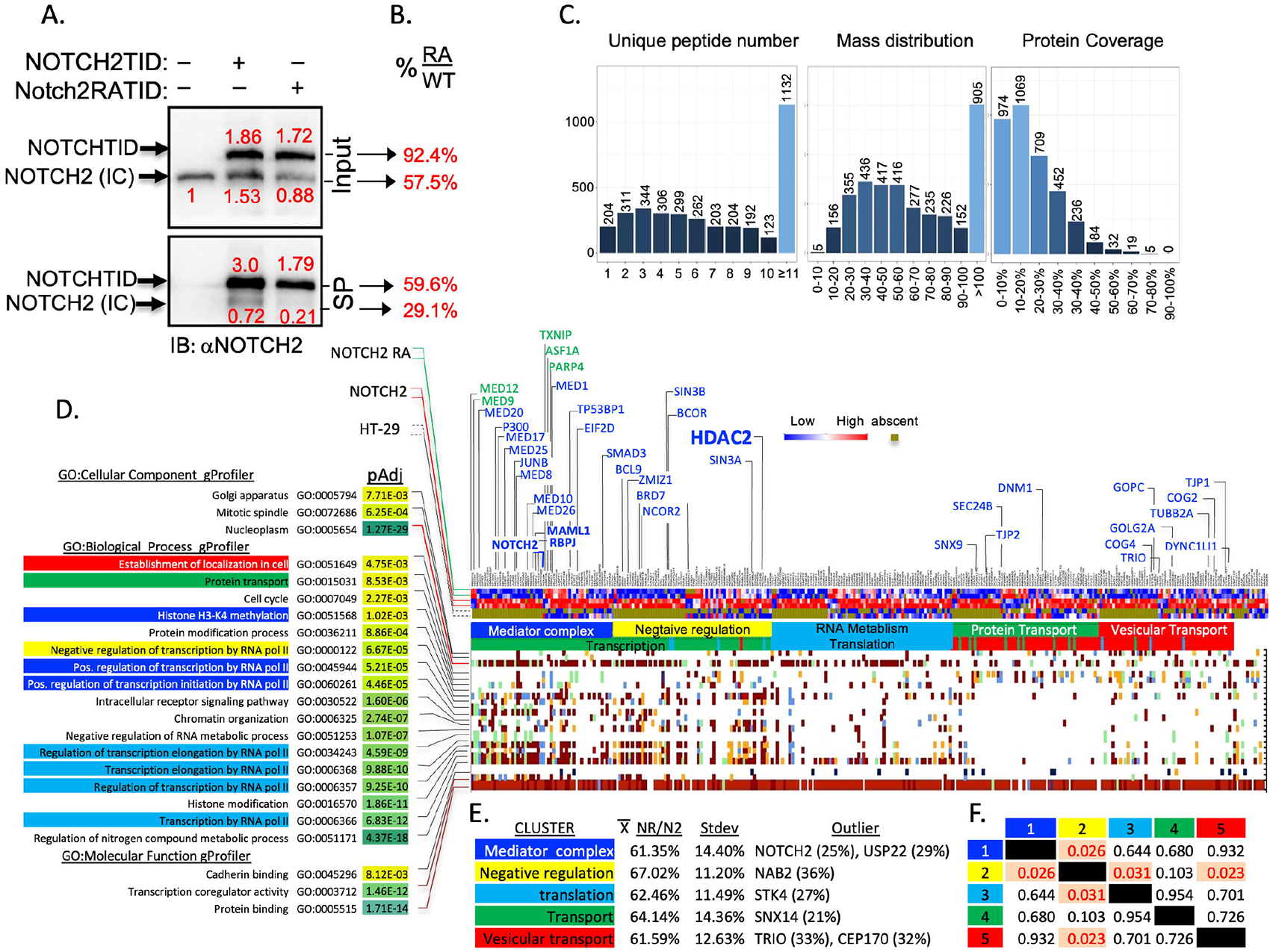
Loss of dimerization dampens protein-protein interactions of NOTCH. **A**. WB analysis of NOTCH2 after biotinylation in NICD2-TurboID and NICD2RA-TurboID stable polyclonal lines. Band intensities were quantified in Image Lab and the amount relative to endogenous NOTCH2 is shown in red above or below the respective band. SP: <u>Streptavidin</u> <u>p</u>ulldown. **B**. The red numerals were used to derive the ratio of the bends in NOTCH2RA sample relative to wild type NOTCH2. **C**. Assessments of data quality including unique peptide number, mass distribution, protein coverage. **D**. ToppFun, gProfiler and String analysis of the biotinylated proteins. The GO term analyses are color coded to match the String cluster shown in E and below the heatmap. Selected proteins are listed above the heatmap, including NOTCH2, RBPj and MAML1 (components of the NTC), HDAC2 (anchor of the yellow cluster) and selected vesicular transport proteins. In blue letters are proteins biotinylated less in NOTCH2RATID; in green are protein biotinylated better by NOTCH2RATID. See Supplemental Table S2 for further detail. **E**. NOTCH2RA biotinylation efficiently relative to NOTCH2 across all clusters, outliers from each group indicated. **F**. Correlation matrix analyses shows that cluster 2 (the HDAC2 anchored cluster) is significantly different in biotinylation efficiency from clusters 1, 3 and 5.

Filtering the data for q_value of >0.05 for the NICD2RA-vs-NICD2 differential analysis revealed that recovery of human NOTCH2 was the most adversely affected by the presence of the RA substitution (Log_2_FC NICD2RA-vs-NICD2; -2.029, or 13% of the NOTCH2 recovered) confirming deficiency in dimer formation.

NOTCH2 is a 2473 amino acids protein and the 17 unique peptide covered 11.4% of it. Aligning all 17 peptides to the NOTCH2 sequence (Q04721) revealed that all peptides mapped to the intracellular domain. To calculate coverage, we performed *in-silico* trypsin digest (https://web.expasy.org/peptide_cutter/) and asked what fraction of possible NICD2 peptides was recovered. This analysis revealed that overall, 32% of the intracellular domain was covered: 42% of the unstructured RAM domain, 18.9% of the ANK domain, and 30% for the PEST domain (S3 Table). To visualize coverage we mapped the biotinylated peptides recovered by MS onto the AlphaFold model of NICD2 and then docked the NICD2 model onto RBPJ/MAML1 structure from [34] using PyMol (https://pymol.org/2/). For docking purposes, the structures were aligned on ANK, and forced the RAM domain of NICD2 to float in space instead of docking on RBPj as it does in the structure. Even occluded residues such K1978 and K2009 were recovered in peptides from TurboID, consistent with robust labeling (Fig S2). These analyses confirm the dimer mutant NICD2RA was nearly ten-fold less effective in interacting with a counterpart, that all interactions were confined to the intracellular domain, and that expected interactors (RBPj, MAML1, MED complex) were indeed recovered.

To further analyze the proteome, we used a cutoff of q_value of >0.1 for the NICD2RA-vs-NICD2, and entered the data into three independent analyses and clustering tools: ToppFun [35], gProfiler [36] and String (see methods for detail). gProfiler established that most of the biotinylated proteins had experimental validation of participation in protein-protein networks (Fig 6D and S2 Table, column AH). Most interactors reside in the nucleus (p=1. 27x10^-29^), the mitotic spindle (p=6.25x10^-04^), or the Golgi apparatus (p=7. 71x10^-03^; Fig 6D and S2 Table columns BF-BH, respectively). String K-means analysis identified 5 significant clusters (Fig 6D and tabs in supplemental Table S2): (1) The blue cluster contained RNA Pol II and positive regulators of transcription, including P300, the core mediator complex, the Set1/MLL complex (H3K4 methyltransferase) and SWI/SNF complex members (facilitator of H2K27 acetylation and nucleosome repositioning). (2) The yellow cluster was anchored around HDAC2 and enriched for co-repressors of transcription and Histone deacetylation complex, including NCOR2 and SIN3A known to antagonize NOTCH signaling. Interestingly, the best studied RBPj co-repressor, SPEN (also known as MINT/SHARP) was not biotinylated by our TurboID constructs (S2 Table). Members of this cluster may interact with RBPj while NOTCH is still in the vicinity, leading to their biotinylation, or are involved in non-histone protein deacetylation [37,38]. (3) Complementing cluster #1, the RNA metabolism cluster (Cyan) contains many proteins involved with co-transcriptional processing of the elongating transcript including the spliceosome complex. (4) The green cluster enriched for transport and receptor mediated endocytosis proteins, and (5), the red cluster enriched in vesicular transport proteins and Golgi proteins (Fig 6D). Importantly, both NICD2 and NICD2RA interacted with each member of these network, however, NICD2RA did so with only 60% efficiency compared to NICD2 across clusters (Fig 6E). Interestingly, cluster #2, the co-repressor cluster, showed the least amount of dimer dependent loss at 67% and was significantly less affected than clusters 1,3 and 5 (Fig 6E and 6F). This may reflect the rate of histone/protein deacetylation near dimeric vs monomeric NTC (or NICD2).

NOTCH proteome analysis in HT-29 colorectal carcinoma cells reveled no dimer specific preferred partner (Fig 6). Monomer preferred partners include ASF1A, a known Notch coactivator [39,40], MED12, MED9, PARP4 and TXNIP (Fig 6D), but since both proteins can exist in complexes as monomers, we did not explore these any further.

### HDAC2 contributes to Notch phenotypes in the intestine

The mild intestinal phenotype may reflect sufficient residual transcription in NDD mice. Alternatively, if the ratio of blue/yellow group interactions reflected higher deacetylation rates near NOTCH2RA bound enhancers, it may have contributed to the reduction in transcription, perhaps by deacetylation of histones and/or other cofactor(s). If HDAC deacetylase activity did not contribute at all, no change would be expected in the presence of HDAC2 inhibitors. If HDAC2 activity synergized with DBZ, HDAC2 inhibition would ameliorate Notch phenotypes. To differentiate between these possibilities, we treated WT small intestine spheroids with Valproic acid (VPA, 2 µM), an HDAC2 inhibitor [41] approved by the FDA for use in humans [42] with the gamma secretase inhibitor DAPT (2 µM), or with both DAPT plus VPA. Inhibition of Notch signaling by DAPT resulted in loss of spheroids proliferation. VPA allowed spheroids to expand at normal rates in the presence of DAPT (Fig 7A). To ask if these effects were seen in vivo, we treated mice with DBZ (10 µmol/kg), VPA (500 or 200mg/kg/day) or DBZ plus VPA. Surprisingly, VPA, which on its own had no measurable effect in this timeframe, preserved the normal morphology of the crypt villus apparatus in the presence of DBZ: goblet metaplasia, Paneth cells differentiation and stem cell proliferation and numbers were all normal relative to the profound changes induced by DBZ alone (Fig 7B and 7C).

**Fig 7.**
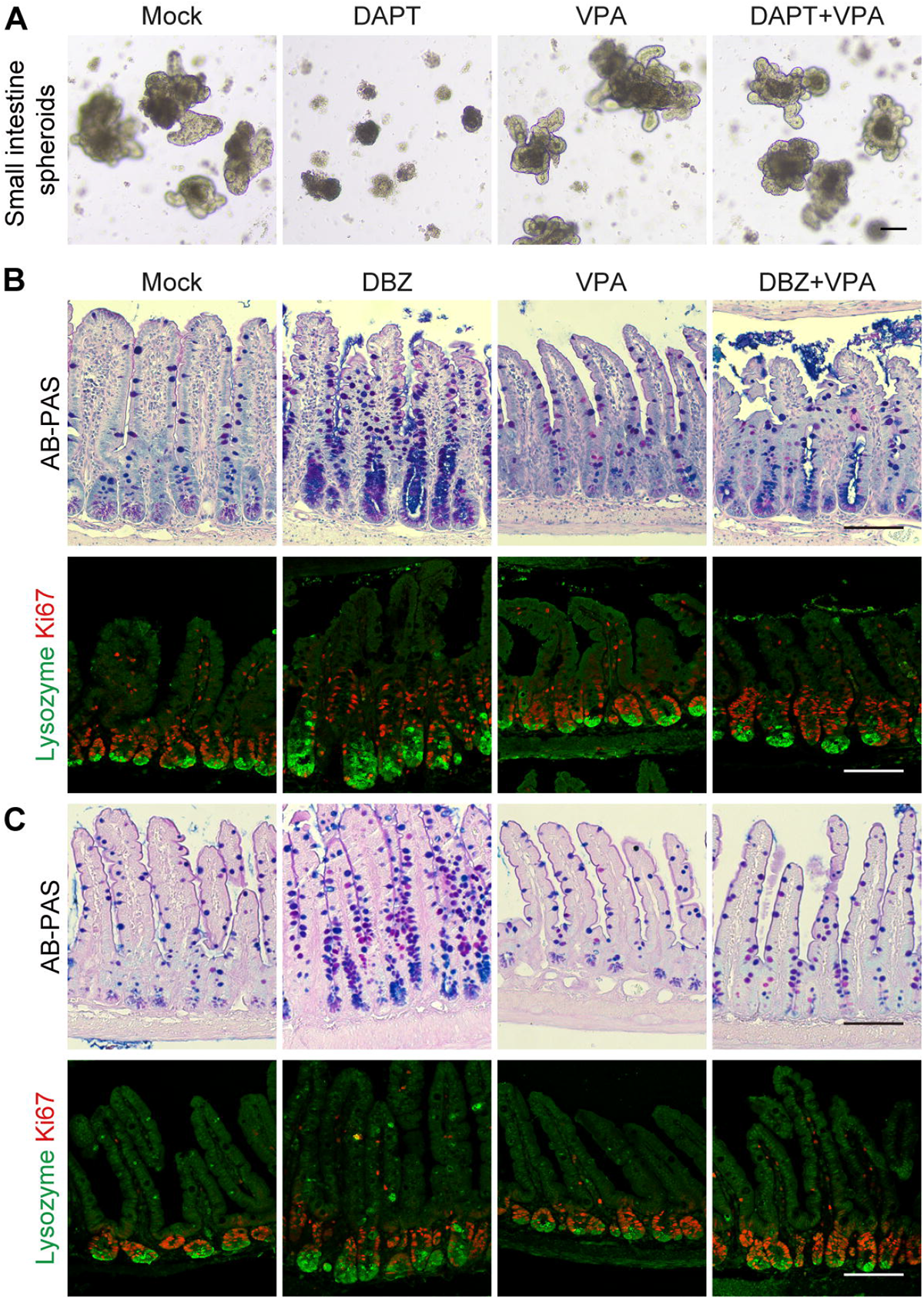
HDAC2 inhibition ameliorates Notch phenotypes in the intestine. **A**. Representative bright-field images from *N1^+/+^; N2^+/+^* small intestine spheroids supplemented with DAPT (2 µM) or VPA (2 µM) alone or DAPT plus VPA. n=3 mice per group. Scale bars=100 μm. **B**. AB-PAS and immunofluorescence staining of Ki67 (red) and Lysozyme (green) in ileum from *N1^+/+^; N2^+/+^* mice treated mice with DBZ (10 µmol/kg) or VPA (500 mg/kg/day) alone or DBZ plus VPA. n=3 mice per group. Scale bars=100 μm. **C**. AB-PAS and immunofluorescence staining of Ki67 (red) and Lysozyme (green) in ileum from *N1^+/+^; N2^+/+^* mice treated mice with DBZ (10 µmol/kg) or VPA (200 mg/kg/day) alone or DBZ plus Valproic acid. n=3 mice per group. Scale bars=100 μm.

## Discussion

### Loss of Notch dimerization sensitizes mice for intestinal injury by elevating the epithelial innate defense against symbionts

NDD mice are more susceptible to DSS-induced stripping of the mucosa and develop colitis when challenged with 1% DSS [14]. In this study we analyzed some of the mechanistic underpinning of this intestinal phenotype. We show that dimerization-deficiency mimics a subset of Notch loss of function (LOF) phenotypes in the small intestine, namely, reduction of stem cells and increase in Paneth cells (Fig 2-4) [20] but no goblet cell metaplasia (S2 Fig, [21,22,24–26]). Mechanistically, our null hypothesis was that monomers-supported activities of NICD will remain unchanged in NDD mice, but dimer-dependent transcription will be affected. Unexpectedly, RNAseq data from colonic spheroids did not identify differential expression of dimer-dependent targets, recording instead overall reduction in Notch-supported function such as proliferation and barrier function (Fig 5, S1 Table). Moreover, spheroids form NDD mice show enrichment for genes associated with host defense against symbionts and interferon-ß targets, consistent with an elevated colonic epithelium innate immunity response in the absence of immune cells or symbionts; none of these transcripts is known to be a direct Notch target. This heightened response to symbionts under basal, laboratory controlled sterile conditions can explain why NDD mice develop colitis when 1% DSS slightly increased exposure to the gut microbiome, or by inference, when fur mites colonized the skin [14].

### Dimer mutation in ICD dampens the vast majority of protein-protein interactions

Since the transcriptomic analysis did not explain why loss of dimerization affected performance in some areas but not others, we investigated the hypothesis the Notch dimers had different proteomic envelop than NICD or monomers, and that the differences might be contributing to the phenotype. We analyzed all interacting partners with TurboID-fused Notch2 and Notch2RA. Notch2 was chosen because it proved critical to the DSS phenotype (Fig 4). Although similar levels of the BirA-fused NICD2 were expressed (Fig 6A), the largest difference observed in recovery was Notch2 itself (Tab S2), confirming that dimers do not form *in vivo*. The experiment revealed that the RA substitution reduced protein-protein-interactions beyond Notch known partners (Fig 6, S2 Table). Interaction with RBPj, MAML1 and P300 were decreased, suggesting lower likelihood of NICD2RA to enter (or stay) in the transcription complex. Indeed, many partners fit into a broad category of positive and negative regulators of transcription, which can be further subdivided into mediator complex and other TFs, negative regulation of transcription, and RNA elongation, splicing, poly adenylation, and translation (Fig 6). Multiple components of mediator complex are less efficiently biotinylated (MED1, MED8, MED10, MED17, MED20, MED25 and MED26). Surprisingly, MED12, part of the cyclin dependent kinase 8 (CDK8) subunit of the MED complex, was a preferred partner of NICD2RA. In *Drosophila*, SPS sites accelerate degradation of Notch complexes by facilitating the recruitment of Med12/Med13/CDK8/cyclinC (CCNC) mediator subunit, leading to CDK8 mediated phosphorylation of the Notch dimer and its degradation [11]. However, MED13 was not differentially biotinylated in HT-29 cells and CDK8 or CCNC peptides were not recovered at all. The significance of preferentially tagging Med12 remains unclear.

The second broad group is involved in transport and is harder to interrogate since dimerization was observed in the context of DNA binding and was not anticipated to impact proteolysis and/or transport. Two clusters of differentially biotinylated proteins are associated with protein transport and endocytosis (cluster 4), and vesicular transport (cluster 5; Fig 6, S2 Table). Endocytosis and intracellular transport are important for Notch localization, activation and regulation [43]. Endocytosis of Notch regulates protein recycling [44] and degradation [45]. Importantly, spatiotemporal analysis using pulses of APEX2 biotinylation [46] mapped discreate Notch interactomes during ligand-induced Notch proteolysis, endocytosis, and nuclear translocation. 696 of 1093 interacting proteins identified by Martin et al were also biotinylated in our experiment (202 of which were nuclear, all shown in S2 Table). Only 85 were differentially labeled by N2 and N2RA, among which were the nuclear proteins Notch2, MAML1, Arid1b, TP53BP1 and Stat1. This suggests to us that intracellular Notch trafficking is largely unaltered by the Arg-Ala substitution, however, the impact of this mutation on trafficking remains to be further analyzed.

As we noted above, NDD mice do not display the full spectrum of Notch LOF intestinal phenotype, specifically, we do not see evidence of goblet expansion (seen even in animals with Notch1 LOF, [25] Fig 3L). Although HDAC3 is a known interactor with co-repressor complex that binds RBPj in absence of NICD1 [47–50], our attention shifted to the HDAC2 cluster. The HDAC2 cluster was significantly less affected by the RA substitution compared to the other clusters. It is possible that Notch2RA/HDAC2 interactions may occur during the “changing of the guard” from active transcription to repression or vice versa [1,51] (Fig4, S2 Table). However, the most important RBPj co-repressor partner, MINT/SHARP/SPEN [52–55] was recovered at similar levels by Streptavidin alone from all cells including parental HT-29 (S2 Table), reflecting low level of basal biotinylation. Because biotinylation of the HDAC2-anchored complex is affected to a lesser degree in Notch2RA-TurboID relative to the reduction in the positive regulator network interactions, we hypothesized that perhaps the HDAC2 interaction may occur when a monomer NTC is sharing SPS with a co repressor complex, or that HDAC2 might play a role outside of transcription, and possibly outside the canonical function of NICD. Regardless, we explored the possibility that deacetylation contributes to the severity of the Notch phenotype.

### HDAC2 inhibition ameliorates the goblet metaplasia seen after NOTCH inhibition in the intestine

Deletion or inhibition of Notch signaling leads to goblet cell hyperplasia and increase in Paneth cells [20,22,24–26]. The relatively modest reduction in HDAC2 interactions may reflect the role of dimerization-mediated cooperativity is to protect against binding of RBPj-corepressor complexes seen in *Drosophila* [56]. Because goblet metaplasia is a dominant phenotype of Notch LOF not recapitulated in weak NotchRA allele, we wondered if HDAC2 synergizes with DBZ. We report that HDAC inhibition can protect the mouse small intestine *in vivo* from the GSI inhibitor DBZ and restore proliferation and Paneth cells numbers in spheroid in *ex vivo* cultures supplemented with DAPT (Fig 7). Amelioration of untoward DBZ effect on the intestine by simultaneous inhibition of HDACs is consistent with data that HDAC inhibitor treatment has ameliorating effect on induced colitis and improves recovery in mice [57–61]. In addition, in an *in vitro* model HDAC inhibition was shown to improve epithelial barrier function [57]. Deletion of HDAC2 alone was shown to protect against DSS [62], however, and in contrast to the benefits of HDAC inhibition, deletion of both HDAC1 and 2 in mice seems to enhance sensitivity to induced colitis and disrupts epithelial barrier function [62–64], indicating that at least some activity is necessary to rip the protective benefits. VPA thus joins Dexamethasone [65] and withaferin A [66] as FDA-approved molecules that alleviate the negative effects of Notch inhibition/loss in the intestine, however, VPA may do so by creating an epigenetic landscape in which the requirement for NICD is lessened.

To conclude, Notch dimerization in mammals seem to be less about transcriptional regulation of an enhancer subset to which they bind through SPS, and more about the overall proteome environment with indirect consequences to innate immunity and barrier functions underlying the hypersensitivity of NDD mice. At present we are not aware of evidence for direct interaction between Notch2 receptors and HDAC2, although HDAC3 is interacting with NICD1 regulating its stability and activity [67]. It is possible that Notch2 and HDAC2 act independently and antagonistically such that co-inhibition has a milder phenotype then the loss of either pathway by reducing or eliminating the need for NICD binding at critical enhancers. Intriguingly, the data we present are also consistent with a model in which NICD or Notch monomers somehow antagonize HDAC2 deacetylase activity, and loss of this activity and subsequent loss of protein acetylation are part of the Notch LOF phenotype. If the putative antagonism reflects direct interaction and non-histone acetylation, it may explain several reports where the canonical model is inconsistent with the data [68,69], and suggest that a subset of acylated proteins are key to the regulation of secretory cell fates. Much remains to be explored.

**Table 1.**
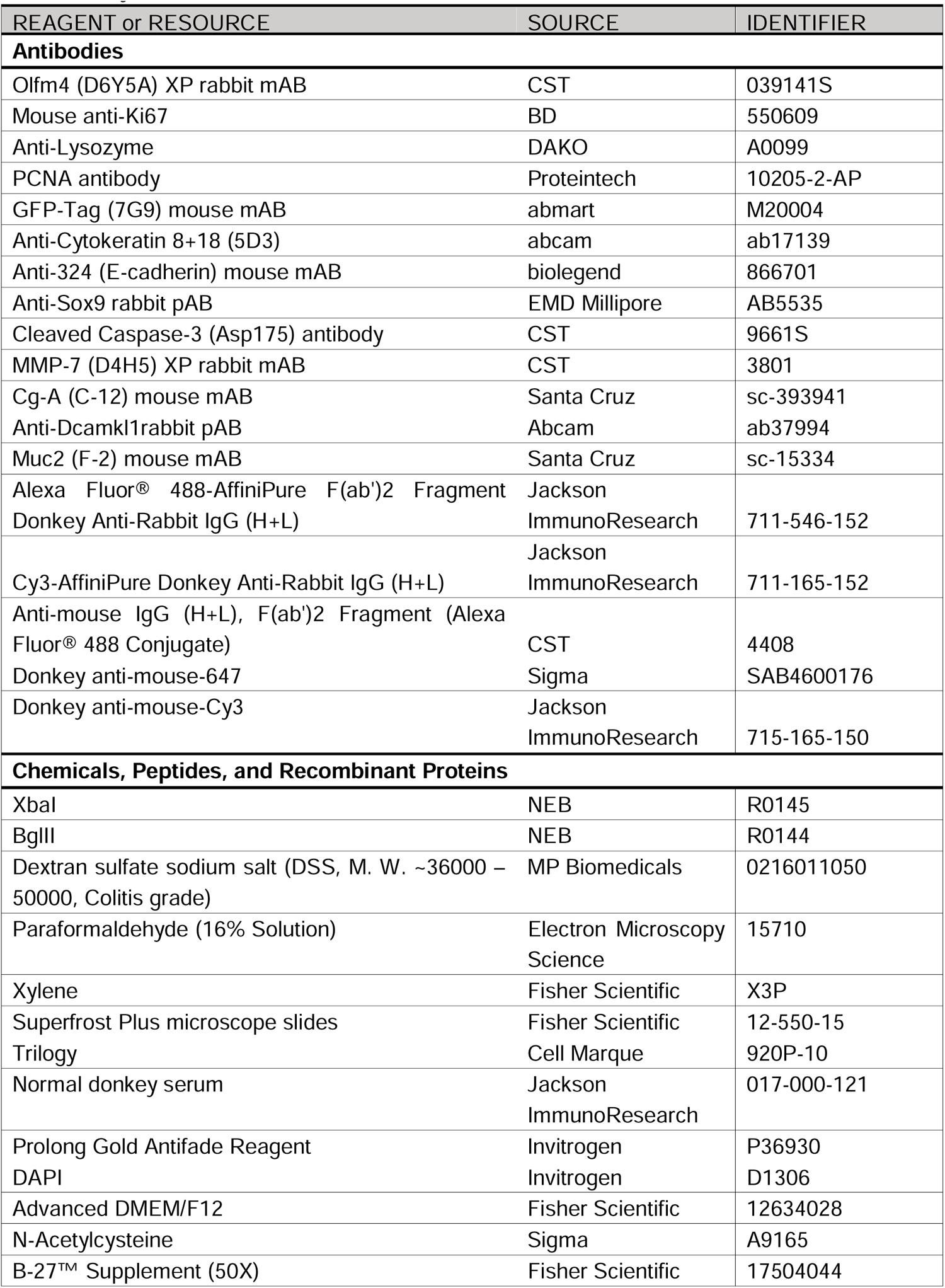
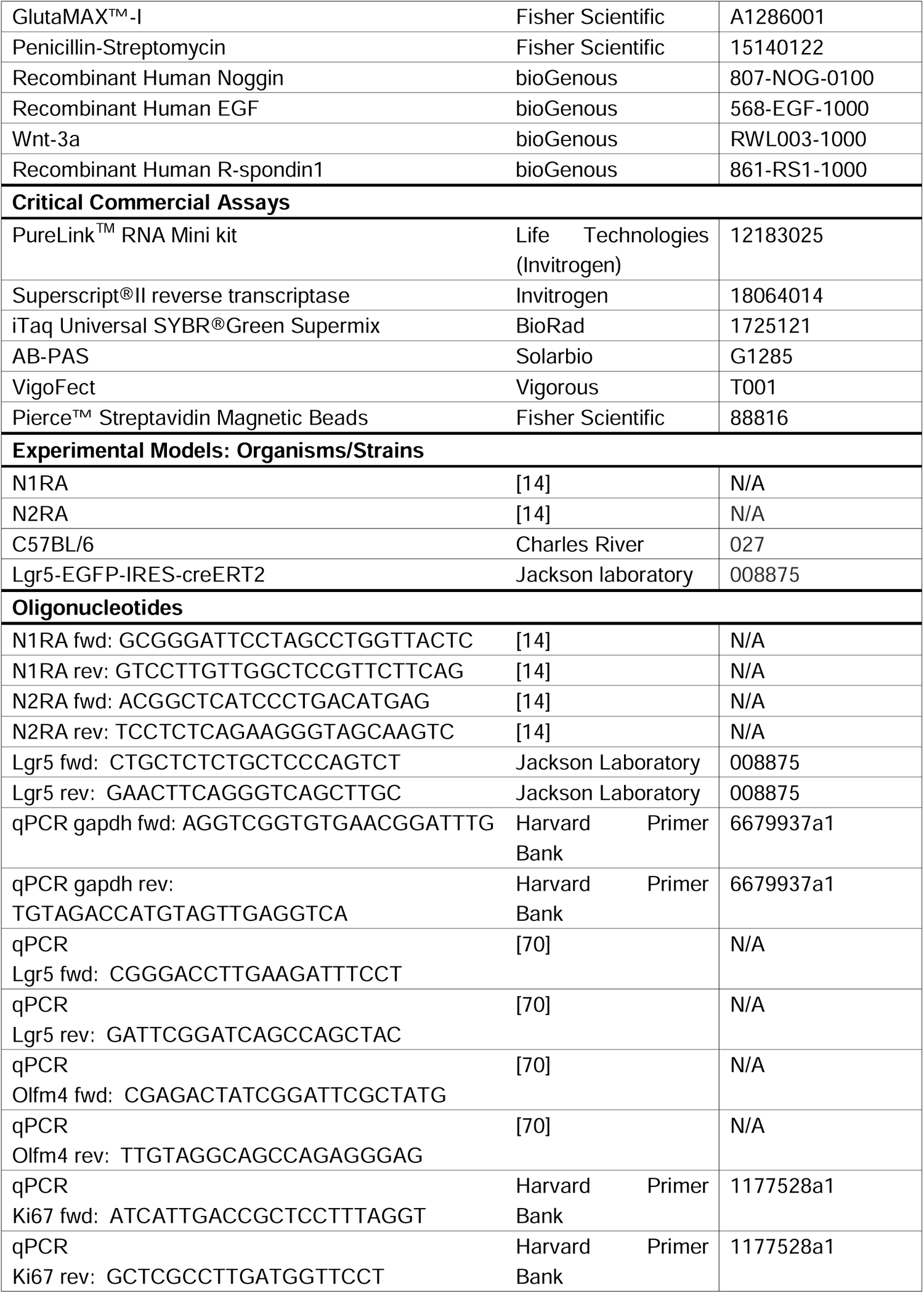
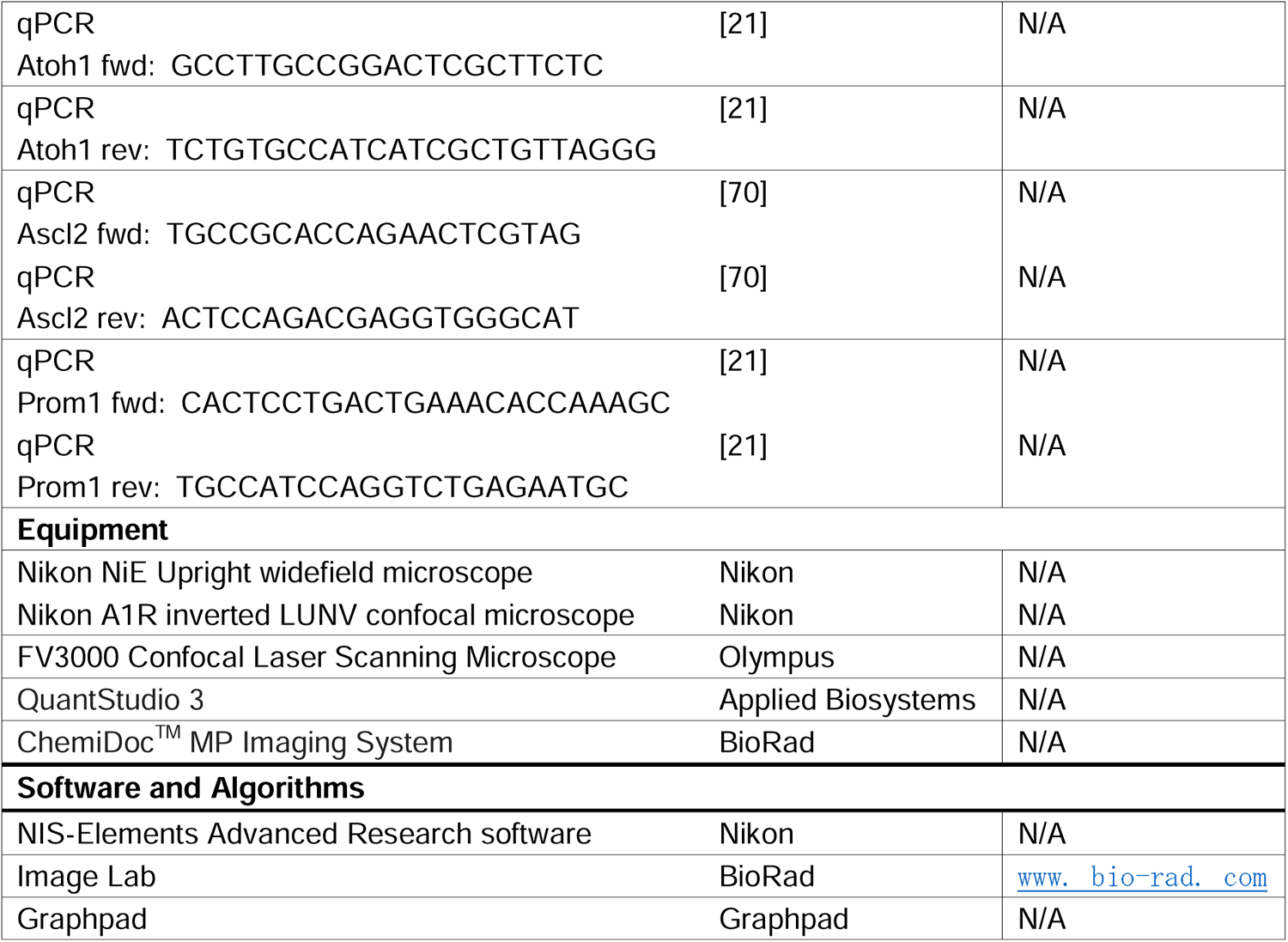
Key Resources/all used materials.

## Materials and Methods

### Ethics statement

All animal studies were approved by the Institutional Animal Care and Use Committee of the Cincinnati Children’s Hospital Medical Center, Ohio, USA (IACUC2018-0105, IACUC2021-0086) and the Institutional Animal Care and Use Committee of Fudan University.

### Animals

All mice were housed at the Animal Facility at Cincinnati Children’s Hospital Medical Center or at Fudan University. Used strains and genotyping information are listed in Key resources table (Table 1).

### Histology

Mouse tissues were fixed overnight in 4% PFA and embedded in paraffin. Adult intestinal tissue was divided into sections corresponding to the jejunum, duodenum, ileum and colon.

Each section was then folded into a Swiss-roll [71] and then fixed, embedded in paraffin and sectioned for histological analysis. 5 µm thick sections were deposited on Superfrost plus microscope slides (Fisher Scientific), deparaffinized with xylene (Fisher Scientific), and rehydrated. Paraffin embedded sections were deparaffinized with xylene and rehydrated. For **immunofluorescence**, the hydrated slides were subjected to Heat-Induced Epitope Retrieval (HIER) by boiling in Trilogy (Cell Marque) for 30 minutes. After cooling, the sections were rinsed with distilled water for five minutes followed by PBS with 0.3% Triton-X100 (Fisher Scientific) for five minutes. The sections were then incubated at room temperature for one hour in blocking solution (10% normal donkey serum (Jackson ImmunoResearch) in PBS-Triton), followed by overnight incubation with primary antibodies at 4°C. The sections were washed 3x10 minutes with PBS-Triton and incubated for one hour at room temperature with respective Fluorophore-labeled secondary antibodies in blocking solution. Unbound secondary antibody was washed off three times for 10 minutes each in PBS-Triton and counterstained with DAPI before mounting with Prolong Gold Antifade reagent (Invitrogen). Confocal imaging was performed on a Nikon Ti-E Inverted Microscope. Where indicated, quantitative analysis of images was done using Nikon’s NIS-Elements Advanced Research software (NIS-AR). For goblet cell and intestinal morphology analysis **AB-PAS** staining (Solarbio, Beijing, China) was performed according to manufacturer’s protocols. In brief, hydrated slides were stained with AB for 15 min followed by washing three times per 2 min with water. The sections were oxidized with periodic acid and washed twice, then stained in Schiff regent for 15 min followed by washing for 10 min. The nuclei were stained by hematoxylin solution for 2 min. Finally, the sections were treated by Scott Bluing Solution for 3 min and washed for 3 min. After dehydrating by series of ethanol and transparent by xylene, the sections were sealed with resinene and determined by blinded pathologists using an optical microscope. For **H&E**, 5µm hydrated sections were stained with hematoxylin for 2. 5 minutes and Eosin for 1 minute before dehydration, clearing, and mounting.

### DSS treatment

To induce colitis mice were given 1% or 2% Dextran sulfate sodium (DSS) in autoclaved drinking water for up to 14 days. Weight was measured every day and stool was checked for changes to monitor disease progression. Mice were either euthanized if severe (>15%) weight loss or blood in stool were observed and tissue collected for analyses. After DSS mice recovered with drinking water w/o DSS for 14 days before starting the next cycle of DSS treatment. At the end of treatment all mice were euthanized, and tissue was collected for analyses.

### qRT-PCR

Total RNA was extracted using the PureLinkTM RNA Mini kit (Invitrogen) according to the manufacturer’s protocols. Purified RNA was reverse-transcribed into cDNAs with SuperScript® II reverse transcriptase (Invitrogen) according to the manufacturer’s protocols. Quantitative Real-Time PCR was performed using iTaq Universal SYBR® Green Supermix (BioRad) and detected on the StepOnePlusTM RT PCR system. Expression levels were normalized to the reference gene Gapdh, data were analyzed using the Delta-Delta-CT methods. A full list of oligos is provided in Key resources table (Table 1).

### Spheroid culture

For small intestine spheroids, jejunal tissue (∼10 cm) was isolated, longitudinally cut, and washed with cold DPBS. Subsequently, the villi were carefully scraped off, and tissue was minced and incubated in DPBS containing 5 mM EDTA for 30 min at 4 °C. After transferring the tissue into a tube with fresh DPBS, crypts were released by vigorously shaking then passed through a 70 μm cell strainer (BD). The isolated crypts were pelleted at 600 rpm for 5 min, resuspended in 25 μL Matrigel (Corning), and seeded in a 24-well plate. After polymerization at 37°C for 15 min, crypts were cultured in the culture medium consisting of advanced DMEM/F12 supplemented with N-Acetylcysteine, B27, GlutaMAX, 1% Penicillin-Streptomycin, 50 ng/mL EGF, 100 ng/mL Noggin, and 500 ng/mL R-spondin1. For colon spheroids, crypts were isolated from the colon and grown as three-dimensional epithelial spheroids in Matrigel as previously described [72]. Colon culture medium contained 50% L-WRN conditioned medium, which was collected from an L cell line engineered to secrete Wnt-3a, R-spondin3, and Noggin [72].

### RNA-seq generation and analysis

Colon spheroids were grown as described above. RNA was isolated from second passage spheroids by using Invitrogen’s Purelink RNA Mini kit according to manufactures directions. Transcriptome libraries and sequencing were performed by the BGI Hongkong Tech Solution NGS Lab to produce over 20 million reads per sample. Raw RNA-seq data was processed using a pipeline developed in-house called CSBB (https://github.com/csbbcompbio/CSBB-v3.0). It employs fastqc to check read quality followed by Bowtie2 + RSEM for alignment and quantification against the mm10 genome. The final outputs are TPM and counts matrices at the gene and isoform level. Differentially expressed genes were then identified using DESeq2 [73]. The DEG were entered into Morpheus (Broad Institute) and unsupervised hierarchical clustering was performed (using Euclidian distance). The kMeans cluster algorithm was deployed to organize the DEG, and putative gene products were further clustered using String (https://string-db.org). Pathway analyses was performed with ToppFun (https://toppgene.cchmc.org/enrichment.jsp) with a B&H FDR cutoff of less than 0.05. The data is deposited at https://www.ncbi.nlm.nih.gov/geo/query/acc.cgi?acc=GSE262485

Reviewer token for GSE262485:

### Lentiviral vector construction and production

For preparation of lentiviruses, NICD2-TurboID, NICD2RA-TurboID and TurboID were cloned into pLVX-P2A-GFP vector. For lentivirus production, HEK 293T cells were transfected in 10 cm plates at ∼60-70% confluency with three plasmids (Core plasmid, psPAX2, and pMD2. G at 7:5:2 ratio) using VigoFect (Vigorous) in culture media according to the manufacturer’s instructions. After 12 hr the media was replaced with 2% FBS-containing media. Approximately 72 h after transfection, the cell medium containing the lentivirus was harvested and filtered through a 0.45 μm filter. To concentrate the virus, the viral supernatant was centrifuged at 200,000 g at 4°C for 2 h and the virus pellet was dissolved in 100 μL advanced DMEM/F12.

### Tissue culture and Turbo-ID

HEK 293T and HT-29 cells were cultured in DMEM (Gibco) supplemented with 10 % FBS and 1% Penicillin-Streptomycin at 37°C in a 5% CO2 environment. For stable HT-29 cell line overexpressing NICD2-TurboID or NICD2RA-TurboID, 40 μL lentivirus was co-incubated with cells in one well of the 6-well plate for two hours at 37°C and shaken every 15 mins. NICD2-TurboID and NICD2RA-TurboID stable polyclonal lines were collected by sorting lentivirus infected cell with GFP. For TurboID, we used culture medium containing a final concentration of 500 μM biotin for one hour. Labeling was stopped after the desired time period by transferring the cells to ice and washing five times with ice-cold PBS. Biotinylated proteins were captured on streptavidin beads according to the manufacturer’s instructions. Purified biotinylated protein samples were submitted to Beijing Genomics Institute (BGI) for next generation label-free quantitative proteomic analyses on a Q-Exactive HF X (Thermo Fisher Scientific, San Jose, CA) in data independent acquisition (DIA) mode.

### Proteomic analysis

50367 peptide and 3580 proteins were quantitated using MSstats software packages. The quantitative statistics of each sample are as follows:

**Table 2:**
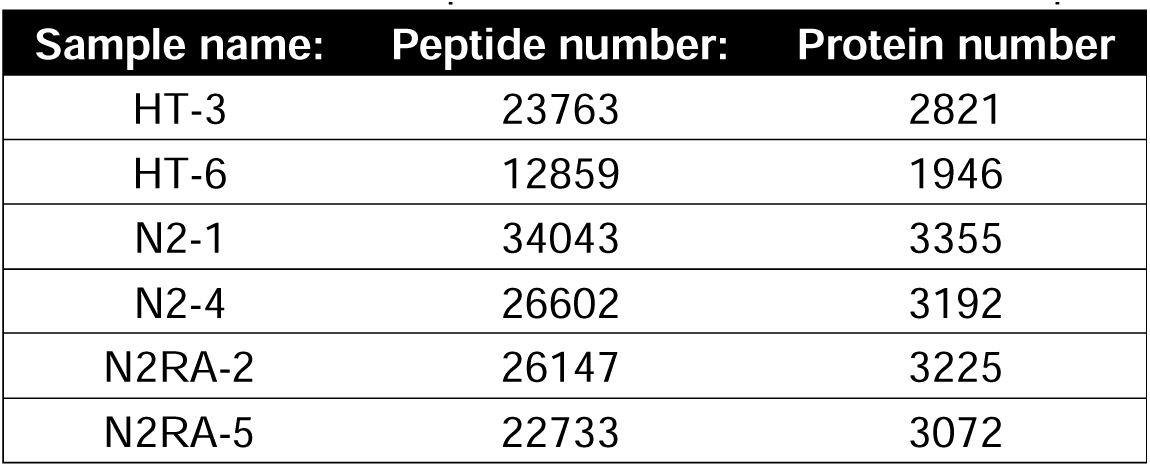
Overview of quantitative results for each sample.

DIA data quality was evaluated based on intra-group coefficient of variation (CV), principal component analysis (PCA), and quantitative correlation of samples. Pearson correlation coefficient of all protein expression between every two samples was represented as a heat map. The CV distribution data, the PCA, and the Heat map of sample correlation analysis are presented in BGI final report (Supplemental Data). To ensure stability and repeatability of the experiment with a large sample size, a quality control (QC) sample comprised of a mixture of all samples was inserted intermittently between the continuous original samples. This allowed experimental conditions to be evaluated by the QC analysis. Initial assessments of data quality, including unique peptide number, mass distribution, and protein coverage indicated the labeling met or exceeded accepted thresholds (Fig 6C and S1 Table). For detailed ethodology consult the BGI Report in Supplemental Data.

### VPA and DBZ treatment of spheroids or mice

For VPA and DAPT treatment of small intestine spheroids, crypts were cultured in medium consisting of advanced DMEM/F12 supplemented with N-Acetylcysteine, B27, GlutaMAX, 1% Penicillin-Streptomycin, 50 ng/mL EGF, 100 ng/mL Noggin, 500 ng/mL R-spondin1 and 2 µM VPA or 2 µM DAPT, or with both DAPT plus VPA. The cell culture medium was changed every other day. For DBZ and VPA treatment of mice, we injected adult mice intraperitoneally daily with 10 µmol/kg DBZ or 200 and 500 mg/kg/day VPA alone, or with DBZ plus VPA (200 and 500 mg/kg/day). The mice were euthanized after X days and their ileum were harvested for histological analyses.

### Quantification and statistical analysis

Statistical analyses were performed using GraphPad Prism software. Two-tailed unpaired Student’s t-tests and a one-way ANOVA were used for comparison between two groups and three or more groups, respectively. Wilcoxon’s rank sum test was used for comparison of body weight. P values <0.05 were considered statistically significant. Bar graphs represent mean ± SEM. All sample numbers (n) listed in legends represent biological replicates.

## Supporting information

S1 Table Transcriptome

S2 Table Proteome

S3 Table Coverage

