## Supplemental Information for "Notch dimerization contributes to maintenance of intestinal homeostasis by a mechanism involving HDAC2"

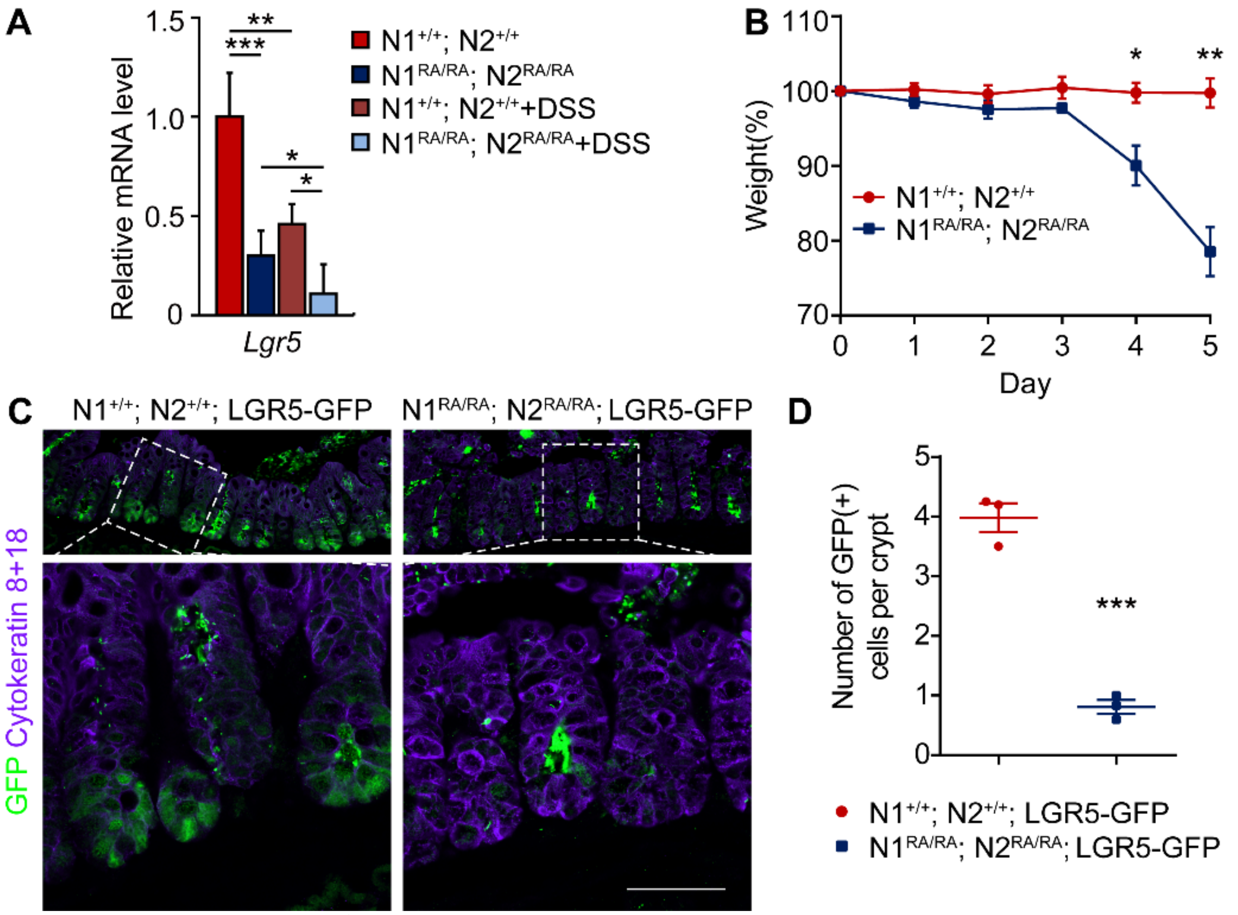


**S1. Fig: NDD mice loose Lgr5$^{+}$ stem cells in the colon.**

A. qPCR of gene expression of *Lgr5* on RNA extracted from distal colon of *N1*$^{+/+}$*; N2*$^{+/+}$ and *N1*$^{RA/RA}$*; N2*$^{RA/RA}$ mice treated for 10 days 1% DSS relative to controls, n=3 mice per group. *$p$ <0.05, **$p$ <0.01, ***$p$ <0.001.

B. Daily weight measurements of *N1*$^{+/+}$*; N2*$^{+/+}$*; Lgr5-EGFP-IRES-creERT2* or *N1*$^{RA/RA}$*; N2*$^{RA/RA}$*; Lgr5-EGFP-IRES-creERT2* mice treated with 2% DSS 5 days, all *N1*$^{RA/RA}$*; N2*$^{RA/RA}$*; Lgr5-EGFP-IRES-creERT2* mice had to be euthanized within 6 days due to severe colitis-induced weight loss. *$p$ <0.05, **$p$ <0.01, ***$p$ <0.001.

C. Immunofluorescence staining of GFP and Cytokeratin 8+18 in distal colon. White dashed box indicated zoomed region. Scale bars=50 µm.

D. Quantification of number of GFP-positive cells per crypt. n=3 mice per group. *$p$ <0.05, **$p$ <0.01, ***$p$ <0.001.

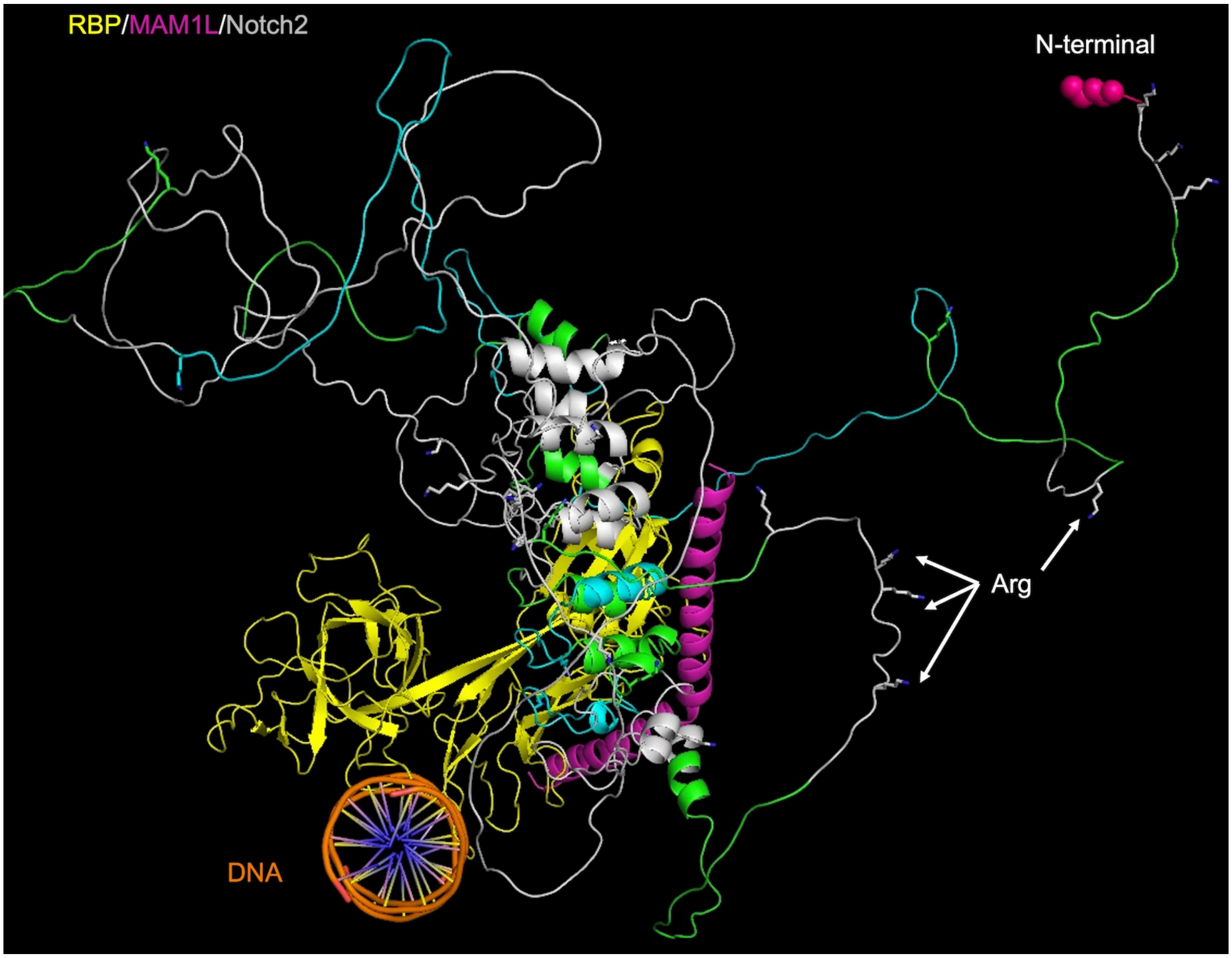


**S2. Fig: Mapping Biotinylated peptide onto NTC.**
Shown are all the biotinylated NICD2 peptides recovered by Streptavidin precipitation using PyMol (https://pymol.org/2/). For docking purposes, the structures were aligned on ANK, such that the RAM domain of NICD2 is floating in space and not docking on RBPj. For NICD2, all lysine residues are shown as “sticks” and the recovered peptides are colored green or cyan to distinguish overlapping peptides. The PEST domain and C-terminus of NICD2 are hidden since modelled it clashed with ANK.

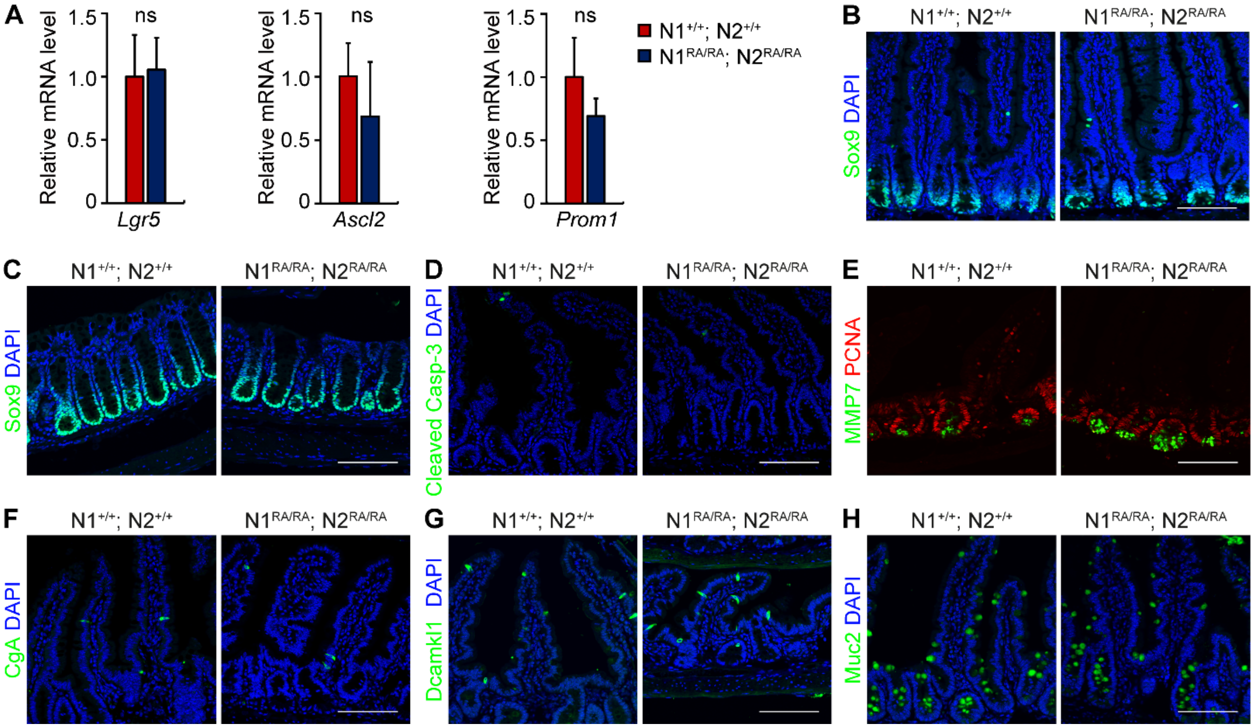


**S3. Fig: Differentiation of intestinal cell types are not affected by Notch dimerization deficiency.**

A. Fold change of *Lgr5*, *Ascl2*, *Prom1* gene expression in jejunum relative to wild type mice analyzed by qRT-PCR. n=3 mice per group. Scale bars=100 µm. Quantitative data are presented as mean ± SEM. ns-Not Significant

B-C. Immunofluorescence staining of Sox9 of jejunum (B) and colon (C).

D. Jejunum immunofluorescence staining of Cleaved Caspase-3.

E. Representative images of MMP7 (Paneth cell marker) and PCNA immunofluorescence in jejunum.

F-H. Representative jejunum immunofluorescence images of CgA (enteroendocrine cells, F), Dcamkl1 (tuft cells, G), and Muc2 (secretory cells, H).

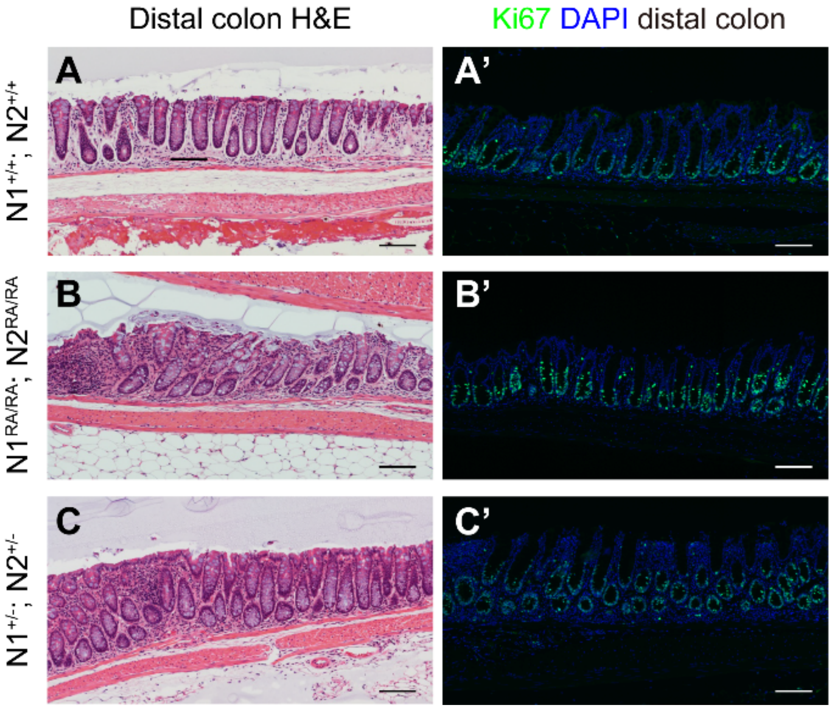


**S4. Fig: Notch dimer-deficiency in mice does not impair colon homeostasis with age.**

A-C. H&E staining of distal colon tissue from aged *N1*$^{+/+}$*; N2*$^{+/+}$, *N1*$^{RA/RA}$*; N2*$^{RA/RA}$ and *N1*$^{+/-}$*; N2*$^{+/-}$ mice. Scale bars=100 µm.

A'-C'. Immunofluorescence staining of Ki67, and nuclei staining with DAPI in distal colon tissue from mice with the indicated genotypes. Scale bars=100 µm. n=3 mice per group.

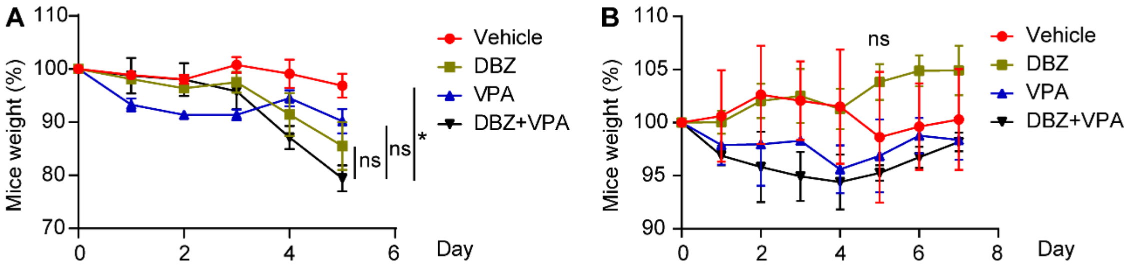


**S5. Fig: Weight loss after five day treatment with GSI inhibitors, VPA, or both combined.**
A. Daily weight measurements of *N1*$^{+/+}$; *N2*$^{+/+}$ mice treated with DBZ (10 µmol/kg) or VPA (500 mg/kg/day) alone or DBZ plus VPA. n=3 mice per group. *$p$ <0.05, ns-not significant.
B. Daily weight measurements of *N1*$^{+/+}$; *N2*$^{+/+}$ mice treated with DBZ (10 µmol/kg) or VPA (200 mg/kg/day) alone or DBZ plus VPA. n=3 mice per group. ns-not significant.

NICD2 amino acid sequence:

SYPLVSVVSESLTPERTQLLYLLAVAVVIILFIILLGVIMAKRKRKHGSLWLPEGFTLRRDASNHKRRE
PVGQDAVGLKNLSVQVSEANLIGTGTSEHWVDDEGPQPKKVKAEDEALLSEEDDPIDRRPWTQQ
HLEAADIRRTPSLALTPPQAEQEVDVLDVNVRGPDGCTPLMLASLRGGSSDLSDEDEDAEDSSAN
IITDLVYQGASLQAQTDRTGEMALHLAARYSRADAAKRLLDAGADANAQDNMGRCPLHAAVAADA
QGVFQILIRNRVTDLDARMNDGTTPLILAARLAVEGMVAELINCQADVNAVDDHGKSALHWAAAVN
NVEATLLLLKNGANRDMQDNKEETPLFLAAREGSYEAAKILLDHFANRDITDHMDRLPRDVARDRM
HHDIVRLLDEYNVTPSPPGTVLTSALSPVICGPNRSFLSLKHTPMGKKSRRPSAKSTMPTSLPNLA
KEAKDAKGSRRKKSLSEKVQLSESSVTLSPVDSLESPHTYVSDTTSSPMITSPGILQASPNPMLAT
AAPPAPVHAQHALSFSNLHEMQPLAHGASTVLPSVSQLLSHHHIVSPGSGSAGSLSRLHPVPVPA
DWMNRMEVNETQYNEMFGMVLAPAEGTHPGIAPQSRPPEGKHITTPREPLPPIVTFQLIPKGSIAQ
PAGAPQPQSTCPPAVAGPLPTMYQIPEMARLPSVAFPTAMMPQQDGQVAQTILPAYHPFPASVGK
YPTPPSQHSYASSNAAERTPSHSGHLQGEHPYLTPSPESPDQWSSSSPHSASDWSDVTTSPTPG
GAGGGQRGPGTHMSEPPHNNMQVYA

NICD2RA amino acid sequence:

SYPLVSVVSESLTPERTQLLYLLAVAVVIILFIILLGVIMAKRKRKHGSLWLPEGFTLRRDASNHKRRE
PVGQDAVGLKNLSVQVSEANLIGTGTSEHWVDDEGPQPKKVKAEDEALLSEEDDPIDRRPWTQQ
HLEAADIRRTPSLALTPPQAEQEVDVLDVNVRGPDGCTPLMLASLRGGSSDLSDEDEDAEDSSAN
IITDLVYQGASLQAQTDRTGEMALHLAARYSRADAAKRLLDAGADANAQDNMGRCPLHAAVAADA
QGVFQILIRNAVTDLDARMNDGTTPLILAARLAVEGMVAELINCQADVNAVDDHGKSALHWAAAVN
NVEATLLLLKNGANRDMQDNKEETPLFLAAREGSYEAAKILLDHFANRDITDHMDRLPRDVARDRM
HHDIVRLLDEYNVTPSPPGTVLTSALSPVICGPNRSFLSLKHTPMGKKSRRPSAKSTMPTSLPNLA
KEAKDAKGSRRKKSLSEKVQLSESSVTLSPVDSLESPHTYVSDTTSSPMITSPGILQASPNPMLAT
AAPPAPVHAQHALSFSNLHEMQPLAHGASTVLPSVSQLLSHHHIVSPGSGSAGSLSRLHPVPVPA
DWMNRMEVNETQYNEMFGMVLAPAEGTHPGIAPQSRPPEGKHITTPREPLPPIVTFQLIPKGSIAQ
PAGAPQPQSTCPPAVAGPLPTMYQIPEMARLPSVAFPTAMMPQQDGQVAQTILPAYHPFPASVGK
YPTPPSQHSYASSNAAERTPSHSGHLQGEHPYLTPSPESPDQWSSSSPHSASDWSDVTTSPTPG
GAGGGQRGPGTHMSEPPHNNMQVYA

**S6. Amino acid sequence of NICD2 and NICD2RA.**
The amino acid sequence of truncated human NICD2 and NICD2RA. The transmembrane domain is boxed in red dashed line. ARG(R1934) and ALA(A1934) are highlighted. Red arrow marks gamma secretase cleavage sites that will generate stable NICD2s.
